# Identifying the most influential features of neural population responses for information encoding and behavior

**DOI:** 10.1101/577379

**Authors:** Ramon Nogueira, Nicole E. Peltier, Akiyuki Anzai, Gregory C. DeAngelis, Julio Martínez-Trujillo, Rubén Moreno-Bote

**Affiliations:** Center for Brain and Cognition & Department of Information and Communication Technologies, Universitat Pompeu Fabra, Barcelona, Spain; Center for Theoretical Neuroscience, Mortimer B. Zuckerman Mind Brain Behavior Institute, Columbia University, New York, New York, USA; Department of Brain and Cognitive Sciences, Center for Visual Science, University of Rochester, Rochester, New York, USA; Robarts Research Institute and Brain and Mind Institute, Schulich School of Medicine and Dentistry, Departments of Psychiatry, Physiology and Pharmacology, Western University, London, Ontario, Canada; Serra Húnter Fellow Programme, Universitat Pompeu Fabra, Barcelona, Spain

**Keywords:** neural coding, decision-making, middle temporal cortex, lateral prefrontal cortex, coarse discrimination, fine discrimination, attention, noise correlations, global modulations

## Abstract

Identifying the features of population responses that are relevant to the amount of information encoded by neuronal populations is a crucial step toward understanding the neural code. Statistical features such as tuning properties, individual and shared response variability, and global activity modulations could all affect the amount of information encoded and modulate behavioral performance. We show that two features in particular affect information: the modulation of population responses across conditions and the projection of the inverse population variability along the modulation axis. We demonstrate that fluctuations of these two quantities are correlated with fluctuations of behavioral performance in various tasks and brain regions. In contrast, fluctuations in mean correlations among neurons and global activity have negligible or inconsistent effects on the amount of information encoded and behavioral performance. Our results are consistent with predictions of a model that optimally decodes population responses, which suggests that in our behavioral tasks the readout of information is near-optimal.

## Introduction

Identifying the statistical features of neuronal population responses that affect the amount of encoded information and behavioral performance is critical for understanding neuronal population coding (Arandia-Romero et al. 2017; Panzeri et al. 2017). The modulation in mean firing rate of individual neurons with respect to a stimulus parameter is a statistical feature that has been typically taken as evidence for encoding information about that stimulus (Hubel and Wiesel 1959; Mountcastle et al. 1967). In addition, changes in network states such as global modulations of activity (Gutnisky et al. 2017; Harris and Thiele 2011; Luczak, Bartho, and Harris 2013), as well as changes in correlated noise among neurons, have been shown to constrain the amount of information encoded by neuronal populations (Ecker et al. 2014; Lin et al. 2015; Schölvinck et al. 2015; Zohary, Shadlen, and Newsome 1994). Indeed, it has been suggested that changes in neuronal tuning, global activity modulations, and noise correlations affect behavioral performance in certain conditions (Cohen and Maunsell 2009; Gu et al. 2011; Mitchell, Sundberg, and Reynolds 2009; Ni et al. 2018). However, the aspects of neuronal responses that most directly affect the amount of encoded information are not clear, since experimental designs often do not allow control over other statistical features that could potentially be involved. Furthermore, it is unknown whether the same features of population responses that affect the amount of encoded information also impact behavioral performance (Arandia-Romero et al. 2017; Panzeri et al. 2017).

In a population of *N* neurons, it is possible to define *N* mean firing rates, *N*(*N*+1)/2 independent covariances, as well as features based on combinations of these quantities. What statistical features matter the most for encoding information? Do these same features also affect behavioral performance? To address these questions, we characterized the amount of encoded information and behavioral performance in three different tasks based on responses of multiple neurons recorded simultaneously in macaque monkeys. We examined neurons recorded in two different brain areas: the middle temporal area (MT), and area 8A in the lateral prefrontal cortex (IPFC). We develop a conditioned bootstrapping approach that allows us to determine the features of neuronal population responses that influence the amount of information encoded and behavioral performance by generating fluctuations of one feature while keeping the other features constant. Using this approach, we found that the amount of information encoded in neuronal ensembles was primarily determined by two features: 1) the length of the vector joining the mean population responses in different experimental conditions (referred here as population signal, PS), and 2) the inverse trial-by-trial variability of the neuronal responses projected onto the direction of the PS vector (projected precision, PP). Contrary to previous suggestions (Ecker et al. 2016; Gutnisky et al. 2017; Kanitscheider, Coen-Cagli, and Pouget 2015; Lin et al. 2015; Zohary et al. 1994), other statistical features, such as mean pairwise correlations (MPC) among neurons and global activity (GA) modulations, did not affect the amount of information encoded when PS and PP were kept constant. Strikingly, we also found that PS and PP are predictive of behavioral performance even though some non-linear processing layers are expected to lie between the information encoding stage and the final stage that generates behavioral choices, whereas MPC and mean GA are not consistently predictive of behavioral performance.

Our results call for a reinterpretation of previous studies that suggested that changes in MPC (Ecker et al. 2010; Gu et al. 2011; Mitchell et al. 2009; Ni et al. 2018; Renart et al. 2010; Zohary et al. 1994) or GA (Ecker et al. 2016; Gutnisky et al. 2017; Kanitscheider et al. 2015; Lin et al. 2015) modulate the amount of information encoded and behavioral performance. Our approach provides a common framework under which all of these results can be reinterpreted by showing that, if PS and PP are held constant, changes in MPC and GA have little effect. Finally, because the same statistical features that modulate encoded information in neuronal populations are the stronger predictors of behavioral performance, our results suggest that the decoding process that drives behavior is optimal or close to optimal.

## Results

We start by showing that fluctuations of MPC and GA influence the amount of information encoded in population activity, consistent with previous studies. We then demonstrate that these effects of MPC and GA are largely eliminated when PS and PP are controlled for, whereas isolated fluctuations of PS and PP strongly predict the amount of information encoded in the population. Finally, we demonstrate that fluctuations of PS and PP are correlated with behavioral performance, and we compare this finding with predictions of an optimal decoding model.

### Mean pairwise correlations and global activity correlate with the amount of encoded information

MPC and GA have been thought to modulate the amount of information encoded in neuronal populations (Ecker et al. 2011, 2016; Gutnisky et al. 2017; Kanitscheider et al. 2015; Lin et al. 2015; Renart et al. 2010; Zohary et al. 1994). We tested the correlation between these statistical features and the amount of information in three different datasets consisting of simultaneously recorded units (2 to ~50 single-units or multiunits) in four monkeys, two brain areas, and three tasks: *i*) pairs of middle temporal (MT) neurons recorded while the animal performed a coarse motion discrimination task (Zohary et al. 1994) (monkey 1, Fig. 1A), *ii*) lateral prefrontal cortex (LPFC, area 8a) neurons recorded with a Utah array while two animals performed an attentional task (Tremblay et al. 2015) (monkeys 2 and 3, Fig 1B), and *iii*) MT neurons recorded with a 24-channel linear array while the animal performed a fine motion discrimination task (monkey 4, Fig. 1C). Behavioral performance in these tasks was defined as the percentage of correctly reported directions of motion (monkeys 1 and 4) or as the mean reaction time when correctly detecting a change in orientation of the attended Gabor patch (monkeys 2 and 3). The amount of encoded information for each dataset was quantified as the cross-validated decoding performance (DP_cv_) of a linear classifier that reads out the activity of the recorded neuronal population to predict *i)* which of two opposite directions of motion was presented in the coarse motion task (monkey 1), *ii)* which Gabor patch was cued in the attention task (monkeys 2 and 3), or *iii)* whether the stimulus motion was right or left of vertical in the fine direction discrimination task (monkey 4). In each case, the classifier was trained on the activity of neuronal ensembles recorded in area MT for the motion tasks or LPFC (area 8a) for the attention task (see STAR Methods for details).

**Figure 1.**
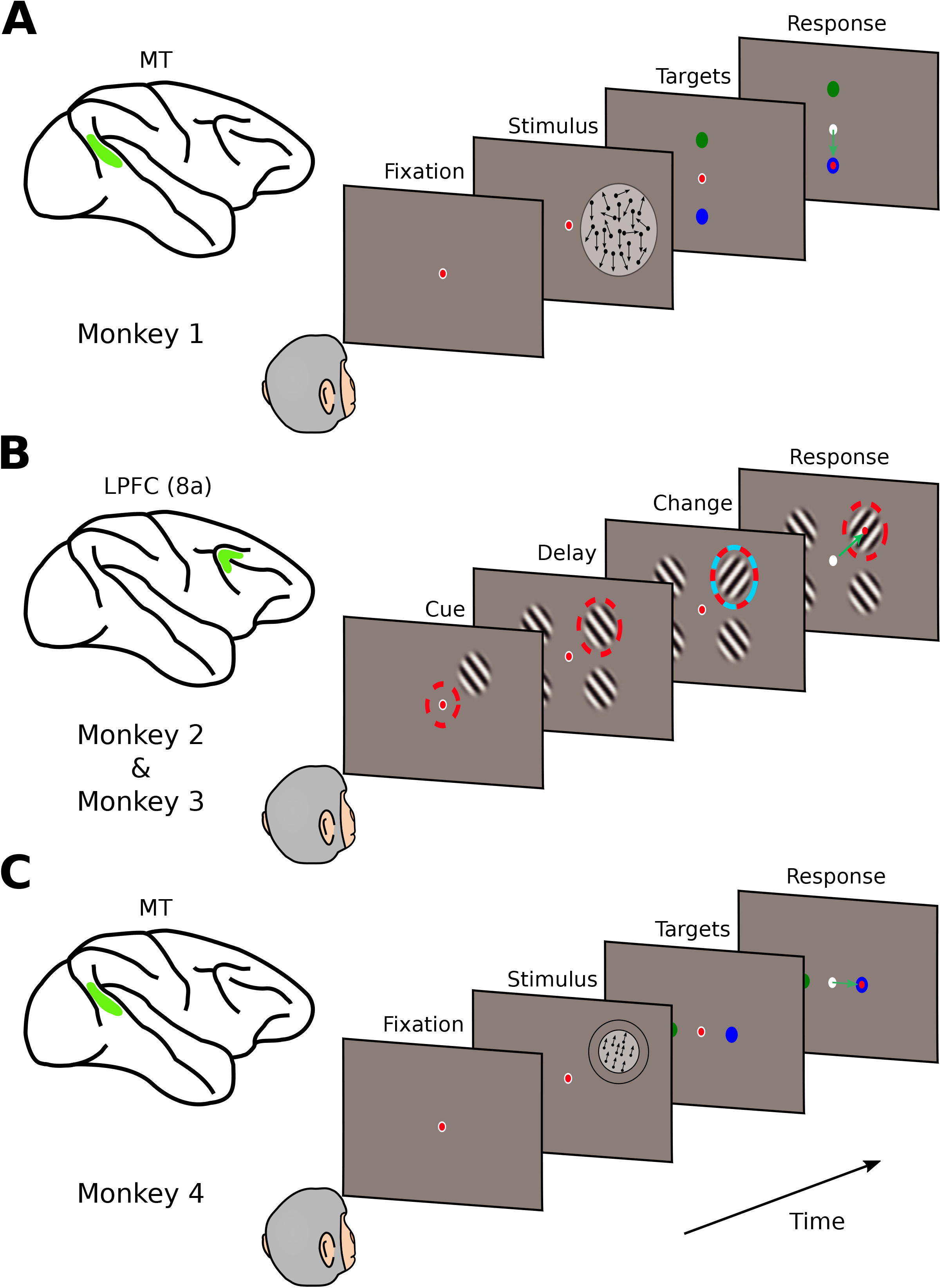
Three behavioral tasks used to test theoretical predictions in macaque monkeys. **(A)** One monkey performed a coarse direction discrimination task (monkey 1) while pairs of units were recorded in the middle temporal (MT) area (Britten et al. 1992). After stimulus presentation (random dots moving toward the preferred or null direction of the neurons under study), the monkey reported the direction of motion by making a saccade to one of two targets. Difficulty was controlled by varying the percentage of coherently moving dots in the stimulus. **(B)** Two monkeys performed an attentional task (monkeys 2 and 3) while ~50 units were recorded simultaneously from the lateral prefrontal cortex (LPFC, area 8a) (Tremblay et al. 2015). Four Gabor patterns were presented on the screen and the task was to make a saccade to the attended location after a change in orientation of the cued Gabor. **(C)** One monkey performed a fine direction discrimination task (monkey 4) while ~25 units were recorded simultaneously from area MT. After presentation of a fully coherent random dot stimulus, the monkey had to report whether dots moved leftward or rightward of vertical by making a saccade to one of two targets. Difficulty was controlled by making the left/right component of motion very small. See STAR Methods for details.

To evaluate whether the amount of encoded information (DP_cv_) was modulated by MPC and GA, we used a non-parametric method to produce fluctuations of MPC and GA generated by bootstrapping trials from each neural recording dataset (Fig. 2A,B). We then examined how fluctuations of MPC and GA affected DP_cv_ (Fig 3; see STAR Methods). We found that an increase in MPC tended to produce a decrease in DP_cv_, therefore reducing the amount of encoded information. This was consistent across all tasks and animals as well as neuronal ensemble sizes. For instance, for monkey 1 (Fig. 3A), an increase of MPC significantly reduced the amount of encoded information by −0.05% (Wilcoxon signed-rank test, *P* = 2.6 10^−3^; see STAR Methods). Qualitatively similar results were found for monkeys 2-4 (Fig. 3B-D).

**Figure 2.**
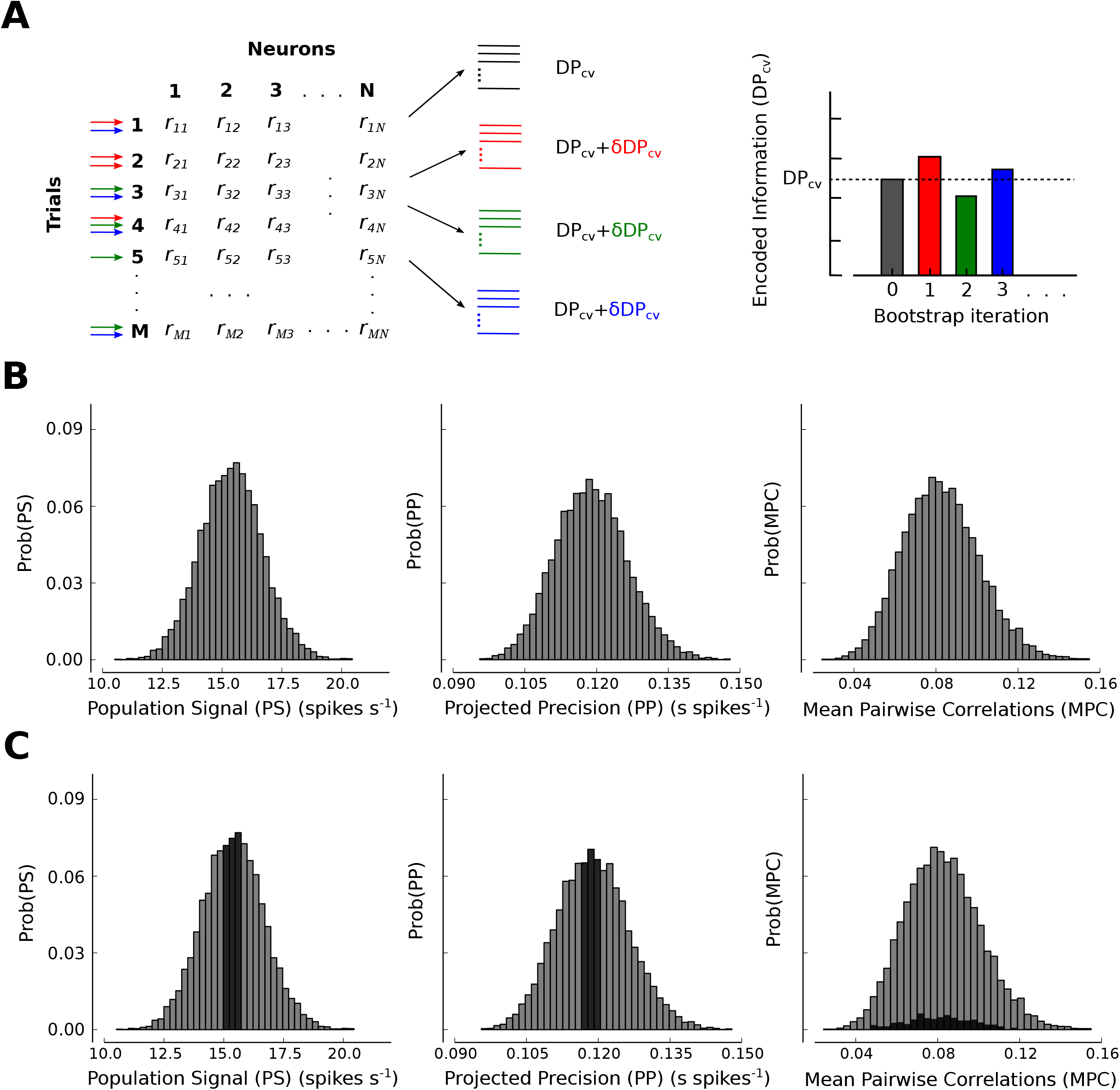
Dependencies between encoded information and statistical features of neuronal responses can be evaluated by the conditioned bootstrapping method. **(A, B)** By subsampling trials with replacement (bootstrap) from the original dataset, distributions of the values of encoded information (DP_cv_) (A) and statistical features of the neuronal activity (B) are generated. Example distributions of the values of population signal (PS), projected precision (PP), and mean pairwise correlations (MPC) for monkey 4 (ensemble size = 10 units). **(C)** A conditioning analysis is performed to determine the impact of fluctuations of each feature of the neuronal response on information by selecting subsets of bootstraps in which the other features are held close to their distributional medians. For instance, to study the effect of MPCs on information, bootstraps are selected such that PS and PP are fixed near their median values (dark regions in the distributions). If bootstrap fluctuations in MPC (while fixing PS and PP) do not modulate DP_cv_, we can conclude that MPC do not play a role in the amount of information encoded by a neural network. This approach is also used to study the effects of bootstrap fluctuations of neuronal activity on behavioral performance.

**Figure 3.**
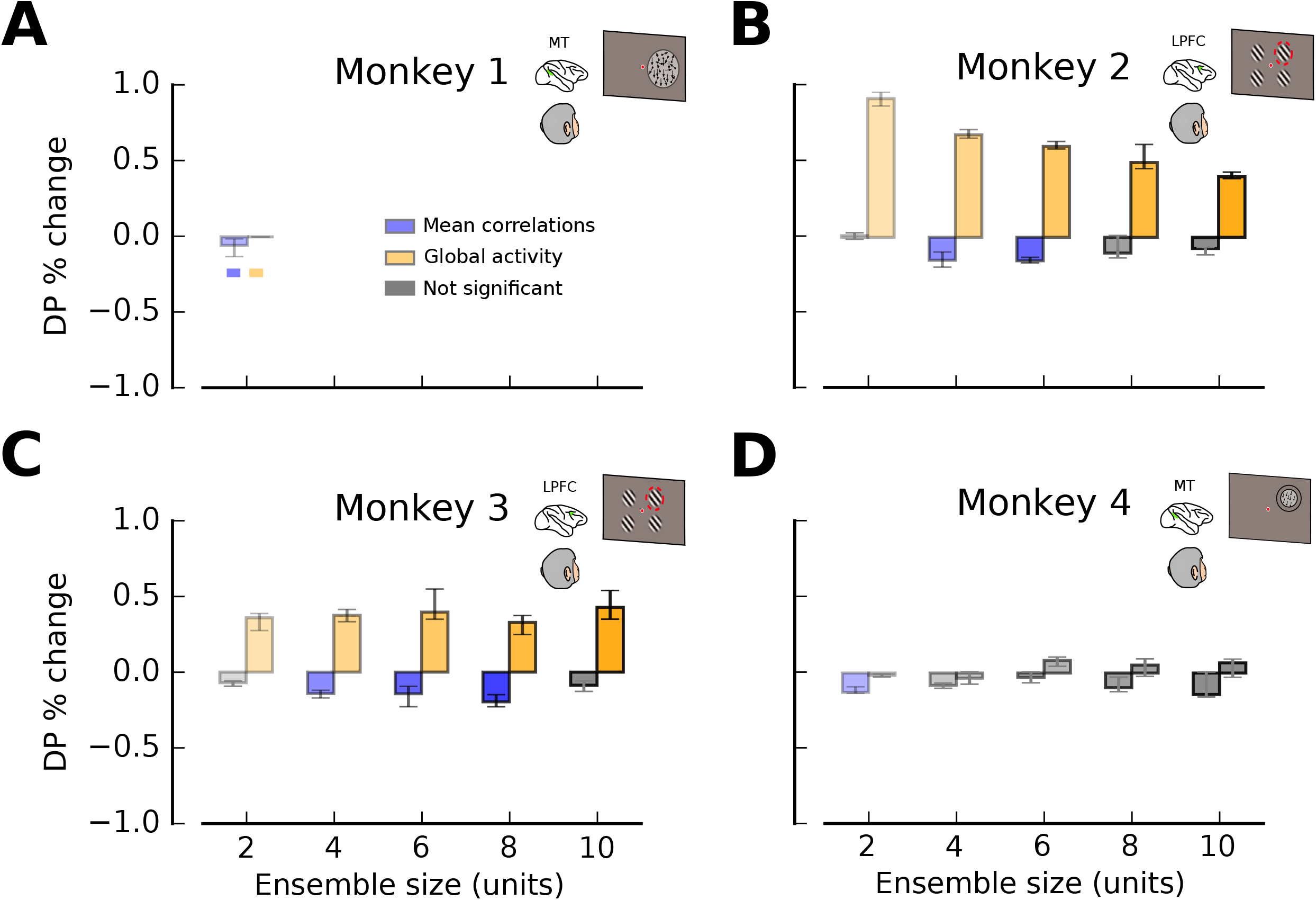
Mean pairwise correlations and global activity correlate with encoded information. **(A)** Percentage change in the amount of encoded information (DP_cv_) related to fluctuations of mean pairwise correlations (MPC; blue) and global activity (GA; orange). Fluctuations of MPC and GA are produced through a bootstrap process in which virtual instances of the same experiment are generated by sampling trials with replacement from a particular dataset (see Fig. 2 and STAR Methods). Smaller values of MPC produce significantly larger values of DP_cv_. Data are shown from pairs of neurons recorded in MT during a coarse direction discrimination task (monkey 1; Figure 1A). **(B, C)** Percentage change in DP_cv_, as a function of ensemble size (2, 4, 6, 8 and 10 units), for recordings from LPFC 8a during an attentional task (monkeys 2 and 3; see Figure 1B and STAR Methods). GA has a strong modulatory effect on encoded information on both monkeys. **(D)** Analogous results for MT ensembles recorded during a fine direction discrimination task (monkey 4; see Figure 1C and STAR Methods). MPC has a consistent negative effect on encoded information. In all panels, error bars correspond to the 25^th^ −75^th^ percentile of the distribution of bootstrap medians and significant deviations from zero are indicated by colored bars (Wilcoxon signed rank test, *P* < 0.05).

In contrast, positive bootstrap fluctuations of GA tended to increase the amount of encoded information for some animals, particularly those from which we made PFC recordings. For monkey 2 (Fig. 3B), bootstrap fluctuations of GA produced significant positive changes in DP_cv_ by 0.91% (ensemble size 2, Wilcoxon signed-rank test, *P* = 6.9 10^−15^; see STAR Methods). For monkey 3 (Fig. 3C), results were qualitatively similar to those obtained for monkey 2, but for monkeys 1 and 4 no significant effect was observed.

Although these results are generally consistent with those from previous experimental and theoretical studies (Ecker et al. 2011, 2016; Gutnisky et al. 2017; Kanitscheider et al. 2015; Lin et al. 2015; Renart et al. 2010; Zohary et al. 1994), the effects are rather small and somewhat inconsistent. As shown below, MPC and GA are confounded with PP and PS and do not exhibit significant correlations with DP_cv_ once PP and PS are held constant.

### Population signal and projected precision determine the amount of information encoded in neuronal population responses

In this section, we identify the statistical features of population tuning and trial-by-trial variability that affect the amount of encoded information (as measured by DP_cv_), and we test our predictions on the neural data described above.

Under some common assumptions (see STAR Methods), an analytical expression for the theoretical decoding performance (DP_th_) of a linear classifier can be derived (Averbeck and Lee 2006) DP_th_ = Φ(*d*′/2), where Φ(*x*) is the cumulative normal function and 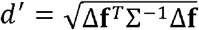 is the signal-to-noise ratio generalized for a population of neurons (Averbeck and Lee 2006; Chen, Geisler, and Seidemann 2006; Gutnisky et al. 2017). The term Δ**f** is the vector joining the means of the population responses in the two stimulus conditions and Σ is the stimulus-invariant noise covariance matrix of the neuronal population. To understand the roles of population tuning and trial-by-trial variability in determining the amount of encoded information, it is useful to re-write this equation by rotating the original *N*-dimensional neural response space along the eigenvectors of the covariance matrix as follows,

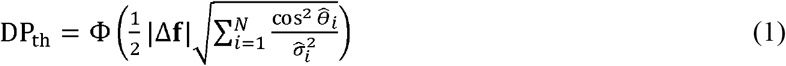

where 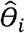 represents the angle between the *i*-th eigenvector of the covariance matrix and the unitary direction defined by the stimulus vector Δ**f**, and 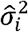 denotes the *i*-th eigenvalue of the covariance matrix (see Fig. S1). Equation (1) reveals that the amount of information encoded by a neural population can be divided into two independent components: the first order contribution from the population tuning (|Δ**f**|; population signal, PS) and the second order contribution from the trial-by-trial variability (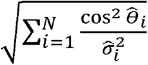; projected precision, PP). It is important to remark that *d*′ is often expressed as the signal to noise ratio of the linear classifier 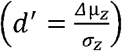, where under optimality, the signal is given by *Δ*μ_*z*_ = Δ**f**^*T*^Σ^−1^Δ**f**, and the noise is given by 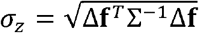. While this *d*′ formulation aims to differentiate signal from noise with respect to the classifier’s decision variable (see Fig. S1; (Averbeck and Lee 2006)), it does not separate the contributions of firing rate modulation (PS) and trial-by-trial variability (PP) of the population to the amount of encoded information, since *Δ*μ_*z*_ and *σ_z_* both contain terms associated with PS and PP.

We find that Eq. (1) provides a very good approximation to the empirical measure of the amount of encoded information, as it accounts for more than 96% of the variance in the cross-validated decoding performance of a linear classifier (DP_cv_) for all datasets (Fig. S2). In addition, Eq. (1) is a better approximation to DP_cv_ than analytical expressions that simplify the covariance structure by removing the off-diagonal terms (Averbeck, Latham, and Pouget 2006; Averbeck and Lee 2006; Pitkow et al. 2015) (see STAR Methods; Fig. S3). Finally, linear discriminant analysis (LDA) produced better matches to DP_cv_ across different ensemble sizes and monkeys than logistic regression (LR) and quadratic discriminant analysis (QDA) did, justifying our use of LDA to evaluate the amount of information encoded by neural populations (DP_cv_; Fig. S4).

Having identified analytically the two features of population tuning and trial-by-trial variability that should determine the amount of encoded information (i.e., PS and PP), we now test the central prediction that information depends exclusively on PS and PP and does not depend on other statistical features, such as MPC and GA, unless those features are correlated with PS and PP. Indeed, we found that bootstrap fluctuations of MPC, GA, PS and PP showed substantial correlations among them (Fig. 4; see STAR Methods). The correlations between fluctuations of PS and GA, PP and MPC, and PP and GA were all highly significant (ensemble size 2, *ρ_PS,GA_* = 0.0073, *P* = 1.2 × 10^−115^; *ρ_PP,MPC_* = −0.061, *P* = 2.4 × 10^−41^; *ρ_PP,GA_* = −0.097, *P* = 10^−136^), whereas the correlation between fluctuations of PS and MPC was not statistically significant (ensemble size 2, *ρ_PS,MPC_* = −0.002, Wilcoxon signed-rank test, *P* = 0.77; see STAR Methods). Therefore, we suspected that the relationships (Fig. 3) between MPC (and GA) and the amount of encoded information would be reduced, if not eliminated, once the modulations in MPC (and GA) were made independent of PS and PP by selecting bootstrap samples with constant PS and PP values.

**Figure 4.**
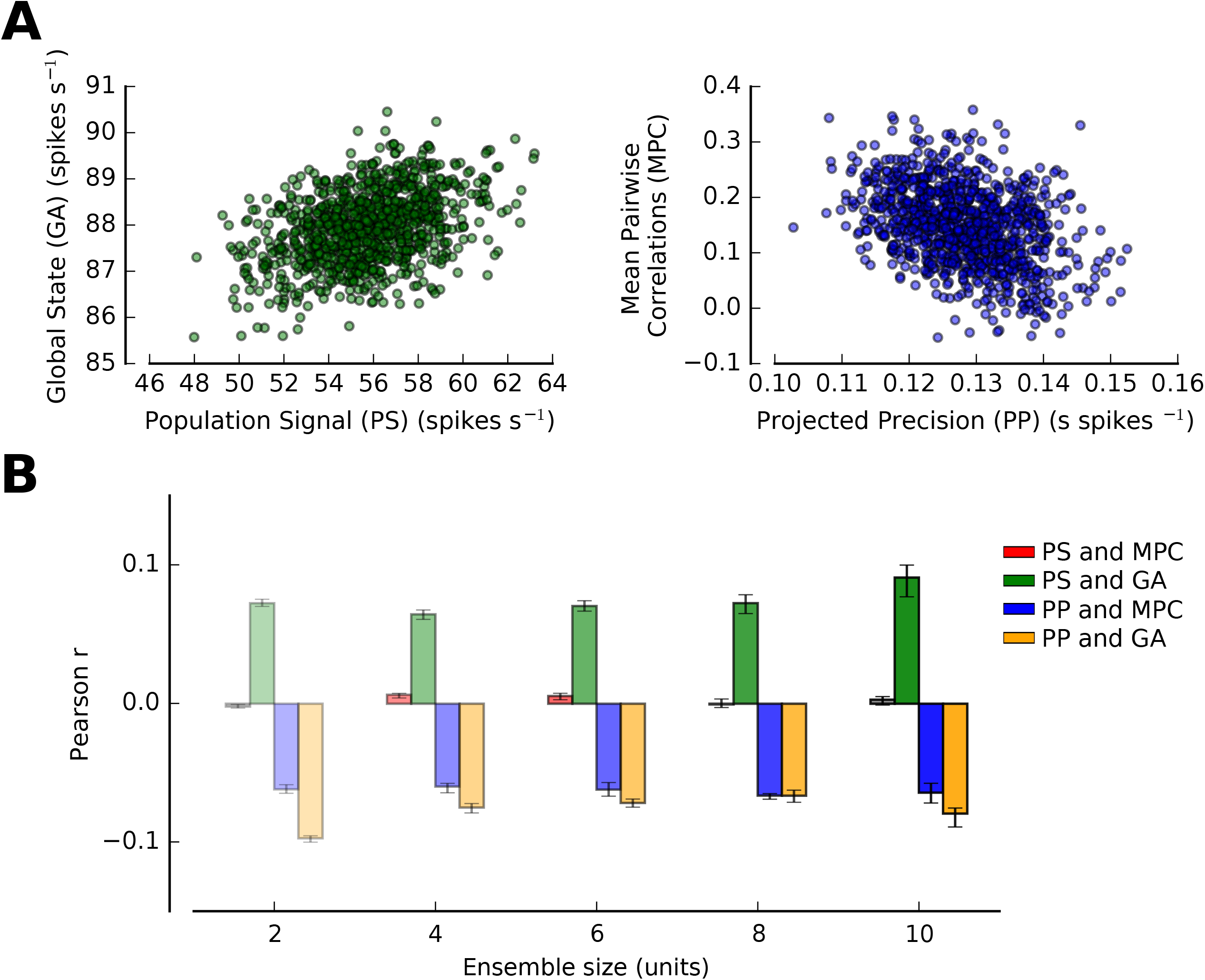
Bootstrap fluctuations in population signal, projected precision, mean pairwise correlations, and global activity are correlated. **(A)** Examples of how bootstrap fluctuations in GA are correlated with bootstrap fluctuations in PS (left panel), and fluctuations of MPC are correlated with PP (right panel) for pairs of neurons recorded in MT during a coarse direction discrimination task (monkey 1; Figure 1A). **(B)** Median Pearson correlation between fluctuations in PS and MPC (red), PS and GA (green), PP and MPC (blue) and PP and GA (orange) across all monkeys for each ensemble size (see STAR Methods). Error bars correspond to the 25^th^ −75^th^ percentile of the distribution of bootstrap medians and significant deviations from zero are indicated by colored bars (Wilcoxon signed rank test, *P* < 0.05). Correlations between these different statistical features of the neuronal activity are likely to explain reported dependencies of encoded information on MPC and GA.

We tested these predictions by using a conditioned bootstrapping method to evaluate the effect of fluctuations in one feature on the amount of encoded information while the values of other features are kept constant (Fig. 2C; STAR Methods). For example, to determine the effects of MPC on DP_cv_ independent of PS and PP, we generated bootstrap samples from the original dataset and selected those bootstrap iterations that produced PS and PP values within a narrow range around their medians. Then, for the selected data, we evaluated the % change in DP_cv_ introduced by the bootstrap fluctuations in MPC. Any dependence of DP_cv_ on MPC, then, cannot be explained by the dependencies of DP_cv_ on PS and PP. This conditioning approach can also be understood as a method for de-correlating different features of the neural activity by keeping only those bootstrap realizations that produced uncorrelated instances of the features to be studied. This method can be used to test the effects of any statistical feature on the amount of encoded information by fixing other features to their representative values, thus isolating the individual effects.

There are other possible approaches to isolating the effects of different statistical features on the amount of encoded information (DP_cv_), such as a model-dependent analysis based on generalized linear models (GLMs). However, due to the linearity assumptions underlying GLMs and the potential collinearity between several statistical features, it is preferable to use a model-independent approach based on conditioned bootstrapping (STAR Methods).

Applying this conditioned bootstrapping approach to our datasets, we found that bootstrap fluctuations of PS and PP greatly affected the amount of encoded information, whereas fluctuations of MPC and GA produced negligible changes in DP_cv_ across different tasks, animals, and neuronal ensemble sizes (Fig. 5). As anticipated, when fluctuations of MPC and GA were conditioned on PS and PP, the effects reported in Fig. 3 largely vanished (Fig. 5, right column). For monkey 1 (Fig. 5A), fluctuations of PS and PP produced significant positive changes in the amount of encoded information by 5.83% (Wilcoxon signed-rank test, *P* = 4.2 10^−37^; see STAR Methods) and 3.21% (*P* = 3.2 10^−38^), respectively. In contrast, fluctuations of MPC and GA had no significant effects on DP_cv_ (*P* = 0.38 and *P* = 0.13, respectively). For monkeys 2-4 (Fig. 5B-D), results were qualitatively similar to those obtained for monkey 1 (see Table S1). We also compared the difference in % change in DP_cv_ for all pairs of statistical features, and found again that PS and PP had the most significant effects on the amount of encoded information (Table S2). The observed dependency of DP_cv_ on PP is generally weaker than that on PS. This could be explained at least partially by the fact that PP is a second order statistic and therefore is expected to be noisier than the first order statistic, PS, when estimated from limited experimental data.

**Figure 5.**
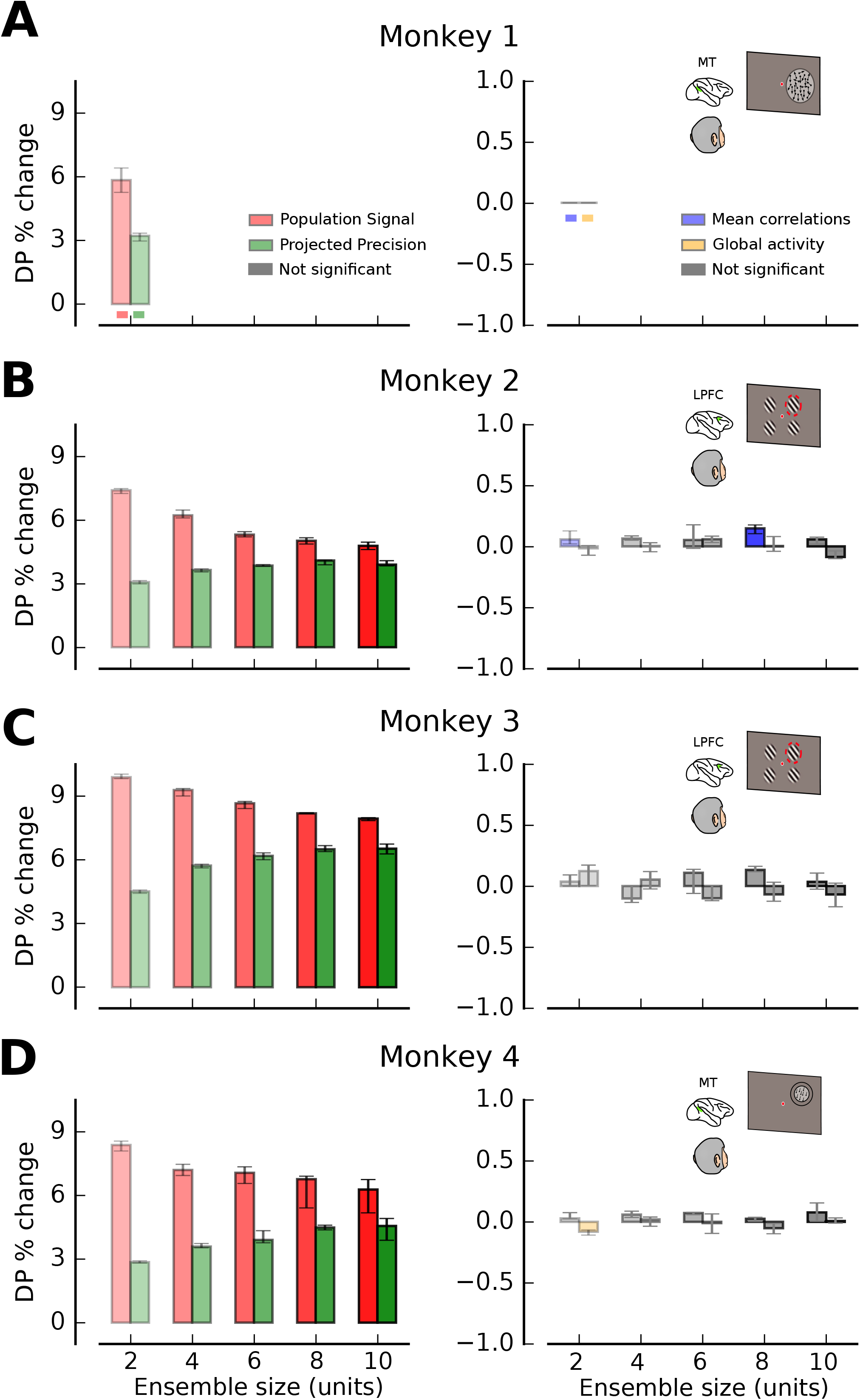
Encoded information depends mainly on population signal and projected precision once other variables are controlled. **(A)** Percentage change in the amount of encoded information (DP_cv_) when changes in one statistical feature of neuronal responses are isolated by the conditioned bootstrapping method (population signal (PS): red; projected precision (PP): green; mean pairwise correlation (MPC): blue; global activity (GA): orange). Only PS and PP produce significant changes in DP_cv_ when other features are keep constant. Data are shown from pairs of neurons recorded in MT during a coarse direction discrimination task (monkey 1; see Figure 1A and STAR Methods). **(B, C)** Percentage change in DP_cv_, as a function of ensemble size (2, 4, 6, 8 and 10 units), for recordings from LPFC 8a during an attentional task (monkeys 2 and 3; see Figure 1B and STAR Methods). **(D)** Analogous results for MT ensembles recorded during a fine direction discrimination task (monkey 4; see Figure 1C and STAR Methods). In all panels, error bars correspond to the 25^th^ −75^th^ percentile of the distribution of bootstrap medians and significant deviations from zero are indicated by colored bars (Wilcoxon signed rank test, *P* < 0.05). See also Fig. S8A.

### Population signal and projected precision are also the strongest predictors of behavioral performance

We have shown that PS and PP are the major statistical features of population activity that affect the amount of encoded information in neuronal populations. But do these features also have an impact on behavioral performance? We reasoned that if downstream neurons that contribute to behavioral choices can extract most of the information encoded by a neuronal population, then behavioral performance should depend on PS and PP while it is largely independent of MPC and GA. However, since there could be many layers of non-linear computations between the information encoding stage and the final behavioral choice, finding such a relationship is not guaranteed *a priori*.

Behavioral performance was measured as the percentage of correctly reported directions of motion (monkeys 1 and 4) or as the mean reaction time (monkeys 2 and 3) (Fig. S5). We first confirmed that modulations in the amount of encoded information (DP_cv_) over different bootstrap samples were significantly correlated with the corresponding changes in behavioral performance (Fig. 6). Although the reported correlations are weak, they are nevertheless consistent with the predictions of a model that reads out neuronal population activity optimally to produce behavioral choices (see next section). We then performed the conditioned bootstrap analysis on behavioral performance separately for each stimulus strength or task difficulty in order to control for trivial dependencies (see STAR Methods). We found that bootstrap fluctuations of PS and PP predicted significant modulations of behavioral performance across different datasets and ensemble sizes, whereas changes in behavioral performance produced by fluctuations in MPC and GA were either weak or inconsistent across different animals and ensemble sizes (Fig. 7). For example, bootstrap fluctuations of PS or PP, while keeping all other statistical features fixed, predicted significant changes in behavioral performance for monkey 1 (Fig. 7A; behavioral change predicted by PS alone:1.41%, Wilcoxon signed-rank test, *P* = 3.3 10^−12^; PP alone: 0.56%, *P* = 2.1 10^−3^). However, the average change in behavioral performance predicted by bootstrap fluctuations of either MPC or GA alone were very small and not statistically significant for this animal (MPC: 0.07%, Wilcoxon signed-rank test, *P* = 0.52; GA: −0.04%, *P* = 0.66). We obtained qualitatively similar results for other monkeys and tasks (Fig. 7B-D; see Table S3), except that bootstrap fluctuations of GA produced significant fluctuations in behavioral performance for monkey 2, but not monkey 3, in the attentional task. We also analyzed the data using Pearson’s correlation coefficient in place of % change in DP_cv_ and obtained qualitatively similar results (Fig. S6). These results were also robust across different conditioning thresholds (Fig. S6; see STAR Methods) and for all pairs of statistical features (Table S4).

**Figure 6.**
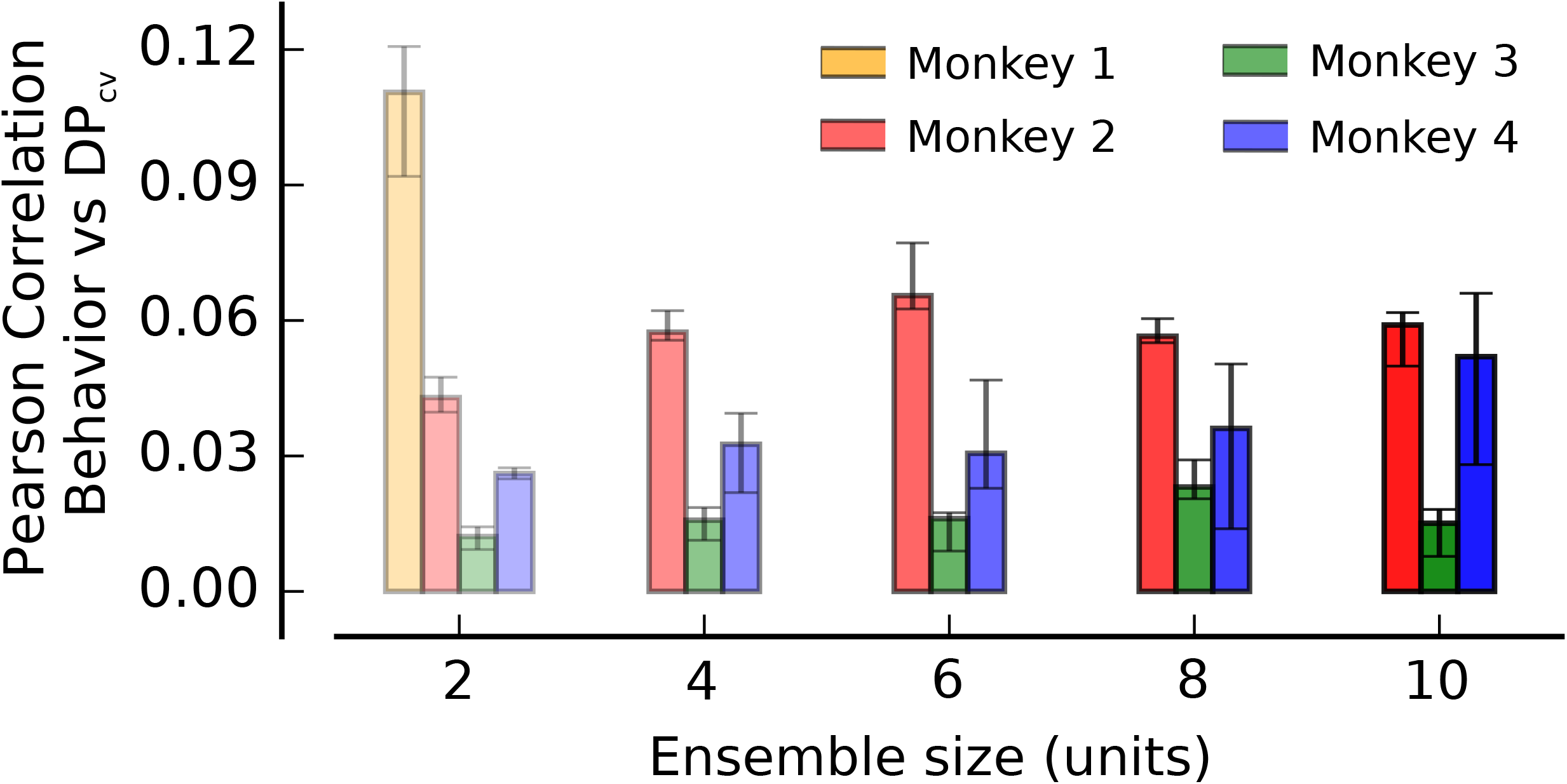
Amount of encoded information correlates with behavioral performance. Pearson correlation between fluctuations in encoded information (DP_cv_) and fluctuations in monkeys performance (see STAR Methods). Across all datasets and ensembles sizes, trials associated with larger encoded information are also significantly associated with better task performance. Error bars correspond to the 25 ^th^ −75^th^ percentile of the distribution of bootstrap medians and significant deviations from zero (colored bars) are calculated by a Wilcoxon signed rank test (not significant if *P* > 0.05).

**Figure 7.**
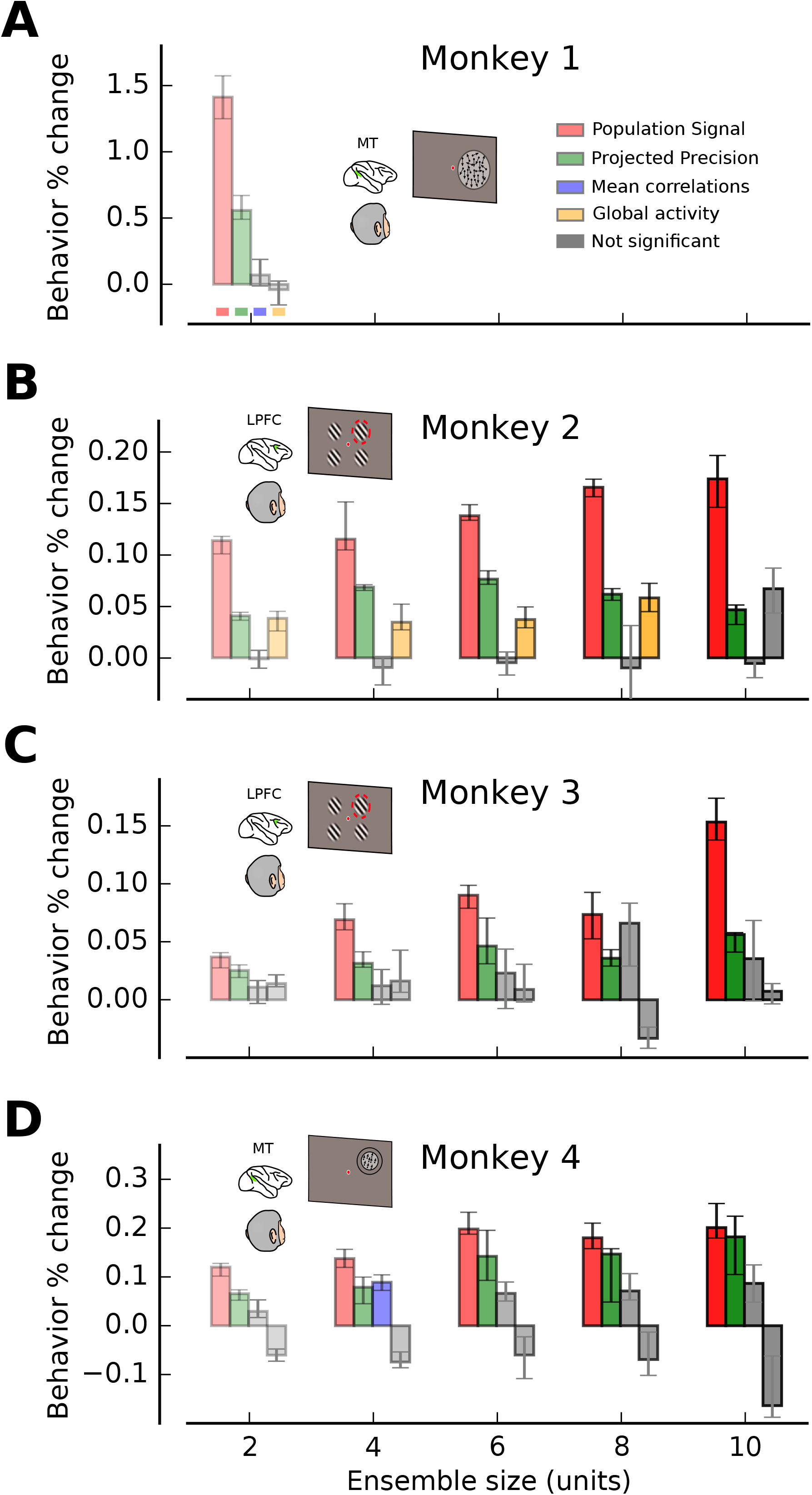
Population signal and projected precision best predict behavioral performance. Percentage change in behavioral performance, as a function of ensemble size, is shown for each behavioral task when the conditioned bootstrapping approach is used to isolate fluctuations in PS (red), PP (green), MPC (blue), and GA (orange). See also Fig. S6 and Fig. S8B.

In summary, we find that PS and PP are the primary statistical features of neural activity that exhibit modulatory effects on behavioral performance, while MPC and GA show little or no consistent effects. These results are in line with those described earlier for the amount of encoded information (Fig. 5). As shown below, the weaker relationship between PS/PP and behavior, as compared to that between PS/PP and the amount of encoded information, is most likely a consequence of behavior being produced as a read-out of the encoding network.

### An experimentally-constrained model accounts for the findings

Although PS and PP are the most consistent and prominent features that correlate with behavioral performance, the observed magnitude of those correlations is quite weak (Fig. 7). One possible explanation for this this weak dependence is that responses are decoded suboptimally, thus producing weak coupling between PS/PP and behavior. Alternatively, the data might be consistent with an optimal readout if behavior is based on much larger neuronal populations than the typical ensemble sizes that we have analyzed. In this latter scenario, PS and PP computed from the entire population of relevant neurons would strongly predict behavior, but PS and PP of small sub-ensembles recorded experimentally would have much weaker correlations with behavior. Indeed, previous work has consistently shown that correlations between neuronal activity of individual neurons and behavioral performance is weak, presumably due in part to the fact that behavior is driven by much larger populations (Britten et al. 1996; Haefner et al. 2013; Nienborg and Cumming 2009; Uka and DeAngelis 2004).

To test whether our experimental findings (Fig. 7) are consistent with an optimal readout of a neuronal population much larger than the observed ensembles, we developed an experimentally-constrained model that could capture the basic statistical properties of the neural responses that we observed (Fig. 8A, Fig. S7A and STAR Methods). The model simulates a number of trials (*M* trials in total) in which a stimulus (*s* = {+1, − 1}) drives a large population consisting of *N* = 1000 neurons. Each neuron has a linear tuning curve with a slope (**m**_1_) drawn from a normal distribution. Neuronal correlations combined limited-range dependencies (Kanitscheider et al. 2015; Kohn et al. 2016), differential correlations(Moreno-Bote et al. 2014) and multiplicative and additive gains (Arandia-Romero et al. 2016). These correlations were generated though a sequence of steps as follows. For each trial *j*, a vector **r**′_*j*_ (size *N* = 1000 neurons), was drawn from a multivariate Gaussian with mean ***μ*** and covariance matrix Σ^gen^. The response vector **r**′_*j*_ was then corrupted with sensory noise, *δs_j_*, which introduces differential correlations and limits the amount of information encoded in large populations of neurons (Moreno-Bote et al. 2014). Multiplicative and additive gains were then applied (Arandia-Romero et al. 2016) (*g_j_*) and a final Poisson step converted each response rate into an observed spike count ***n***_*j*_ (Kanitscheider et al. 2015) (see STAR Methods). Fano factors (FF) that roughly matched typical values found experimentally were used (Arandia-Romero et al. 2016; Nogueira, Lawrie, and Moreno-Bote 2018; Shadlen and Newsome 1998). Finally, based on the neuronal activity generated by the network, we generated surrogate choices (*c_j_*) by optimally decoding the whole population of 1000 simulated neurons 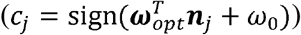 on a trial-by-trial basis. The presence of differential correlations ensures that the amount of encoded information could not scale indefinitely with the number of neurons.

**Figure 8.**
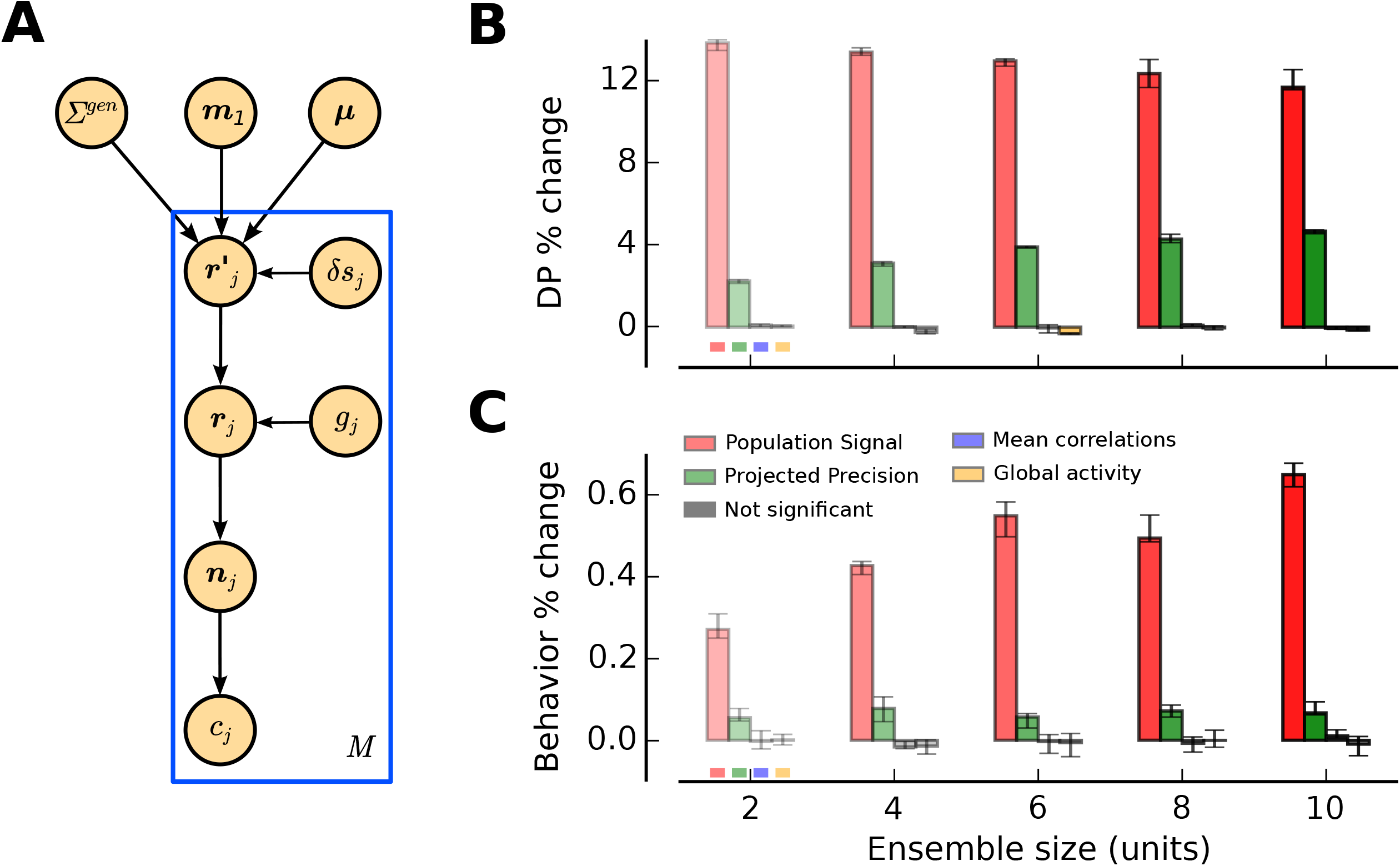
An experimentally-constrained neural population model accounts for the empirical findings. **(A)** Generative model for the simulated neuronal responses. A population of *N* model neurons was characterized by linear tuning curves with slope **m**_1_. An intermediate activity pattern 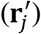 was obtained by drawing an *N*-dimensional sample from a multivariate Gaussian distribution (mean ***μ*** and covariance *Σ^gen^*) and then corrupting it with sensory noise (*δ_s_j__*) on every trial *j* (**r**_*j*_) (Mtrials in total). A homogeneous response gain modulation (*g_j_*) and a Poisson step were applied to produce the final population spike count (***n***_*j*_). The choice of the virtual agent was obtained by an optimal read-out of the population activity pattern on each trial (STAR Methods). **(B)** Percentage change in the amount of information encoded by the model (DP_cv_), as a function of the ensemble size, when bootstrap fluctuations of different features of the neural activity are isolated. Only isolated bootstrap fluctuations in PS and PP influence the amount of information encoded by the population, consistent with the experimental observations (Fig. 5). **(C)** Percentage change in behavioral performance predicted by fluctuations in different statistical features of model population activity. PS and PP are the most important factors influencing behavioral performance. The magnitude of changes in behavioral performance is similar to that observed experimentally (Figs. 6 and 7). See also Fig. S7.

To compare to our experimental data, we then randomly selected small ensembles (up to 10 units) out of the full population of 1000 model neurons to test how fluctuations in PS and PP of the small ensembles correlate with encoded information and simulated behavior of the full network (Fig. S7), repeating the analyses performed on the experimental data (Figs. 5 and 7). Consistent with the experimental data, PS and PP computed from these small ensembles of model neurons are the only features that correlated significantly with DP_cv_ and surrogate behavioral performance, while MPC and GA did not (Fig. 8B,C). We also found that changes in the model’s simulated behavioral performance associated with bootstrap fluctuations of PS and PP were quite small (Fig. 8C; ensemble size 10; surrogate behavioral change predicted by PS alone: 0.63%, Wilcoxon signed-rank test, *P* = 1.73 10^−6^; PP alone: 0.065%, *P* = 0.028) and roughly similar in magnitude to the experimental results (Fig. 7). As expected, when neuronal ensembles of up to 100 neurons were used instead, fluctuations of PS and PP had a larger effect on surrogate behavioral performance (ensemble size 100; surrogate behavioral change predicted by PS alone: 0.90%, Wilcoxon signed-rank test, *P* = 1.73 10^−6^; PP alone: 0.21%, *P* = 4.38 10^−3^). Finally, correlations between fluctuations of DP_cv_ and simulated behavioral performance were weak (Fig. S7G) and comparable to those observed in the experimental data (Fig. 6). Therefore, the weak correlations that we observed experimentally between PS/PP and behavioral performance are broadly consistent with a scenario in which behavior is generated by an optimal read out of a much larger neuronal population of neurons.

## Discussion

Identifying the statistical features of neural population responses that determine the amount of encoded information and predict behavioral performance is essential for understanding the link between neuronal activity and behavior. Based on data collected from two brain areas and three behavioral tasks, we identified two critical features: the length of the vector joining the mean responses of the population of neurons across two conditions to be distinguished (population signal, PS), and the projection of the inverse covariance matrix onto the direction of that vector (projected precision, PP). We found that changes in PS and PP are significantly correlated with the amount of encoded information and behavioral performance, but MPC and GA are not, once their correlations with PS and PP are eliminated. Our experimental results are consistent with predictions of a neuronal population model with realistic neuronal tuning, variability, and correlations that is read-out optimally to generate behavioral choices.

Although previous studies have examined how input signals are represented in the average activity of neurons (Hubel and Wiesel 1959; Mountcastle et al. 1967; Renart and van Rossum 2012) and have described the types of neuronal correlations that can benefit or harm encoded information (Abbott and Dayan 1999; Averbeck et al. 2006; Ecker et al. 2011; Moreno-Bote et al. 2014; Zohary et al. 1994), they have not clearly identified the primary features of neuronal population activity that constrain the amount of encoded information and affect behavioral performance. As shown above, a traditional interpretation of d’ considers the signal and noise components of the classifier’s decision variable. However, it does not separate the contributions of population tuning (first order statistic) and trial-by-trial variability (second order statistic) of the neuronal responses to the amount of encoded information. Thus, one important contribution of our study is identifying PS and PP as the critical features of neuronal activity that modulate the amount of encoded information and predict behavioral performance. Another contribution of this study involves demonstrating that other features of neuronal activity, namely MPC and GA, have little or no impact on the amount of encoded information and behavioral performance once PS and PP are controlled for, even though they have previously been considered important factors (Arandia-Romero et al. 2016; Ecker et al. 2016; Gutnisky et al. 2017; Kanitscheider et al. 2015; Lin et al. 2015; McAdams and Maunsell 1999; Shadlen and Newsome 1998; Zohary et al. 1994). We developed a novel approach for studying correlations among multiple variables while eliminating the effects of other variables (conditioned bootstrapping method). This method uses bootstrapping to generate continuous distributions of multiple statistical features so that fluctuations of some features can be computed for a subset of bootstrap samples that yielded no fluctuations in other features. This method allowed us to isolate the effects of a particular feature on both the amount of encoded information and behavioral performance. Thus, our approach clearly differs from the traditional trial shuffling method that destroys all dependencies in the data to examine the effects of correlations (Kohn and Smith 2005; Leavitt et al. 2017; Romo et al. 2003; Tremblay et al. 2015). It also differs from maximum entropy models that generate surrogate data while fixing the first and second moments of the generated distribution to desired values (Elsayed and Cunningham 2017).

Previous studies have suggested that MPC and GA affect behavioral performance (Cohen and Maunsell 2009; Gu et al. 2011; Mitchell et al. 2009; Ni et al. 2018). On the surface, these studies appear to be at odds with our main finding that only PS and PP should affect the amount of encoded information or behavioral performance. Indeed, we find that bootstrap fluctuations of MPC and GA have little effect on the amount of encoded information or behavioral performance when PS and PP are kept constant. Our results suggest that the previous studies found effects of MPC and GA on behavioral performance because these variables are themselves correlated with PS and PP (see Fig. 4). In fact, there may be many other statistical features of neuronal responses that seemingly influence the amount of encoded information and behavioral performance, but such relationships could be explained as a byproduct of correlations of these statistical features with PS and PP.

Although theoretical research has proposed that fluctuations of GA can modulate the amount of encoded information (Ecker et al. 2016; Kanitscheider et al. 2015; Lin et al. 2015), a recent study based on large neuronal populations in monkey primary visual cortex found that the amount of information does not vary with large fluctuations in GA (Arandia-Romero et al. 2016). Another recent study found a modest, but significant, increase in decoding performance when population activity decreased (Gutnisky et al. 2017), which appears to be inconsistent with our theoretical prediction. Again, however, some of the previous reports of positive effects of GA may have arisen from correlations between fluctuations in GA and PS. Indeed, we have found a highly significant correlation between PS and GA. Therefore, an important question is whether there is a residual effect of GA on the amount of encoded information in the previous studies after the contributions of PS and PP are eliminated. Our theory predicts that there should not be, and the conditioned bootstrapping approach that we have developed would be useful for teasing apart the statistical features (such as PS and PP) that fundamentally affect the amount of encoded information and behavioral performance from heuristic parameters such as GA that may have only secondary effects on information or behavior.

In conclusion, based on information metrics that are readily applicable to neuronal data, we developed a theoretically-driven analysis that has identified the statistical features of neural population responses that modulate the amount of information encoded by cortical neuronal populations. We found an excellent agreement between the theory and the experimental results, which indicates that the assumptions of our approach are reasonable. A critical finding is that PS and PP are the primary features that correlate with behavioral performance and that one must be careful when interpreting effects of other statistical features that may co-vary with PS or PP. Finally, the fact that the same features affect both the amount of encoded information and behavioral performance suggests that the decoding process that drives the animal’s behavior is optimal or close to optimal. Indeed, if the decoding process were strongly sub-optimal, the most relevant statistical features for encoding information would not match those affecting behavior; rather, the relevant statistical features would take different forms depending on the exact set of read-out weights used by downstream areas to produce behavior.

## Acknowledgements

We thank K.H. Britten, J.A. Movshon, W.T. Newsome, M.N. Shadlen and E. Zohary for making their data available in the ‘Neural Signal Archive’ (http://www.neuralsignal.org/), and W. Bair for maintaining this database. We thank Matthew Leavitt for useful comments on the manuscript. R.N. is supported by NSF’s NeuroNex program (DBI-1707398) and by Gatsby Charitable Foundation (GAT3419). R.M-B. is supported by BFU2017-85936-P and FLAGERA-PCIN-2015-162-C02-02 from MINECO (Spain) and the Howard Hughes Medical Institute (HHMI; ref 55008742). G.C.D., N.E.P., and A.A., as well as experiments in the DeAngelis lab, were supported by NEI grants EY013644 and EY016178.

## Author Contributions

R.N., J.M.T., G.C.D. and R.M.B. conceived the project. R.N. and R.M.B. derived the analytical calculations. R.N. performed the analysis. N.E.P., A.A., and J.M.T. obtained the recordings (monkeys 2, 3 and 4). All authors wrote the manuscript.

## STAR Methods

### Contact for reagent and resource sharing

Further information and requests for resources and reagents should be directed to and will be fulfilled by the Lead Contact, Ramon Nogueira

### Experimental model and subject details

See next section

## Methods details

### 1. Theoretical expression for the amount of encoded information

#### 1.1. Theoretical decoding performance for an arbitrary linear classifier

If we assume that the activity of a neuronal population **r** follows a multivariate Gaussian distribution, the covariance matrix for **r** is stimulus-independent, the probability of presenting condition 1 is same as presenting condition 2 (*p*(*C*_1_) = *p*(*C*_2_) = 0.5), and the classification is based on a linear projection of **r** into a scalar variable *z* = ***ω***^T^**r** + ω_0_ (linear classifier) (see (Fisher 1936)), then the performance of the linear classifier can be expressed as

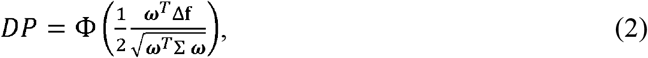

where Δ**f** ≠ ***μ***_1_ − ***μ***_2_, ***μ***_1_ = *E*[**r** | *C*_1_, *C*_1_], ***μ***_2_ = *E*[**r** | *C*_2_], Σ = *E*[(**r** − ***μ***_1_)(**r** − ***μ***_1_)^*T*^ | *C*_1_] = *E*(**r** − ***μ***_2_)(**r** − ***μ***_2_)^*T*^ | *C*_2_], Φ(·) is the cumulative Gaussian function, and *DP* is the decoding performance. This expression gives the percentage of correct classifications (i.e. *DP*) that would be achieved by a linear classifier that reads out from a neuronal ensemble using an arbitrary set of weights, ***ω***.

#### 1.2. Theoretical DP for the optimal linear classifier

By optimizing Eq. (2) with respect to ***ω***, we find that ***ω***_*opt*_ ∝ Σ^−1^ Δ**f**, which corresponds to the solution for the Linear Discriminant Analysis (LDA)(Fisher 1936). We can substitute ***ω*** for ***ω***_*opt*_ in Eq. (2) to find the DP for the optimal classifier

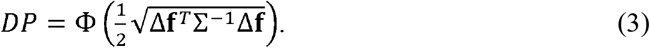

The term inside Φ(·) is known as 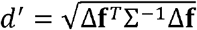 (Averbeck and Lee 2006; Chen et al. 2006). It is a scalar quantity and therefore it remains invariant under unitary rotations of the reference frame. By rotating the original neuron-based orthogonal basis so that Σ (and thus Σ^−1^) becomes diagonal, we can express Eq. (3) as

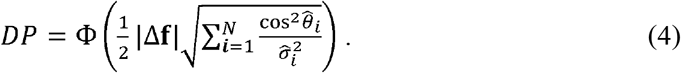

(see (Moreno-Bote et al. 2014) for a similar derivation in the case of linear Fisher information). The first term, |Δ**f**| which we will refer to as Population Signal (PS), is the norm of the stimulus tuning vector Δ**f**. It measures the overall modulation of the activity of the neuronal population as a function of the stimulus conditions. The second term, 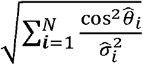 which we will call Projected Precision (PP), is a function of 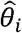, the angle between the *i*-th eigenvector of the covariance matrix Σ and the direction of the stimulus tuning vector Δ**f**, and 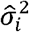, the *i*-th eigenvalue of the covariance matrix. Thus, Projected Precision is the square root of the sum of squares of the precision of the population activity along each of the ellipsoid’s N axes (N = the number of neurons) projected on to the axis of the normalized stimulus tuning vector 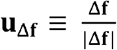. This rotation allows us to dissect the independent contributions of population tuning (first-order statistics) and trial-by-trial variability (second-order statistics) to the amount of information encoded by a neural population, which is not possible with the standard factorization of d’ into signal and noise components (see main text).

#### 1.3. Theoretical DP for shuffled neuronal recording data

Trial shuffling is a commonly used technique to destroy the trial-by-trial shared fluctuations among neurons while preserving their mean firing rate to different experimental conditions. To understand the effects of noise correlations on DP, it is useful to derive theoretical expressions of DP for shuffled data. Shuffling the activity of single neurons across trials for a fixed stimulus condition transforms the covariance matrix Σ into Σ_*sh*_ (Averbeck et al. 2006), which is only approximately diagonal due to finite data size effects. The theoretical expression of DP for the shuffled data is given by

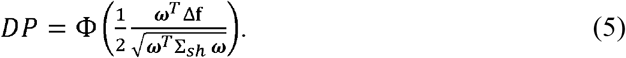

The optimal classifier is therefore 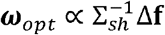, and Eq. (4) becomes

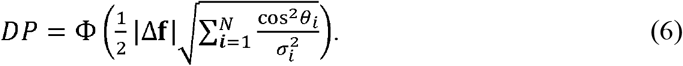

In this expression, PS remains the same as in Eq. (4) while the PP for the shuffled data becomes 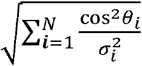, where 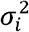 is the variance of the response of each neuron and *θ_i_* is the angle between the stimulus tuning axis **u**_Δ**f**_ and each vector of the neuron-based orthogonal basis defining the original N-dimensional space of the neuronal activity. The above equation can also be expressed as

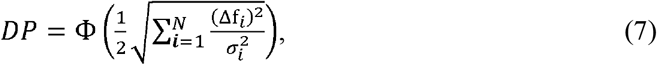

where 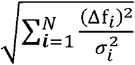 can be understood as the square root of the sum of the signal-to-noise ratio (SNR) of all neurons in the ensemble (Seung and Sompolinsky 1993).

#### 1.4. Theoretical DP under suboptimal read-outs

Here, we derive theoretical expressions of DP for two sub-optimal classifiers: one blind to response variability and another blind to pairwise correlations (Pitkow et al. 2015).

The variability-blind classifier takes into account the neuronal tuning but it is blind to any elements of the covariance matrix (considers the covariance matrix to be the identity matrix). Thus, the readout weights for this classifier are given by ***ω*** ∝ Δ**f**. Introducing this expression into Eq. (2), we find

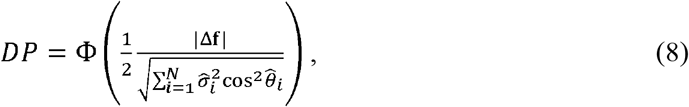

where 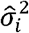 and 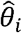 are as defined before.

The correlation-blind classifier (Pitkow et al. 2015) only takes into account the signal-to-noise ratio of each neuron, and ignores pairwise correlations among the neuronal ensemble (off-diagonal terms in the covariance matrix). The set of weights for this classifier, then, is given by 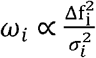, which can also be written as 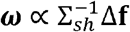. By substituting this expression into Eq. (2), we obtain

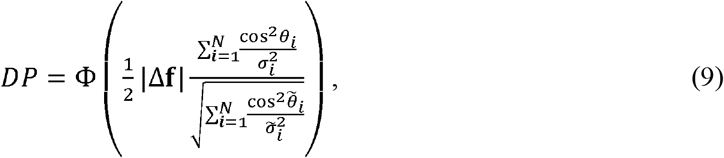

where 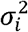 and *θ_i_* are as defined earlier and 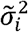 and 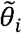 correspond to the *i*-th eigenvalue and the angle between **u**_Δ**f**_ and the *i*-th eigenvector of the matrix 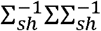, respectively. The correlation blind classifier, though suboptimal for correlation-intact data, is optimal for shuffled data from which pairwise correlations have been removed.

#### 1.5 Theoretical expression for DP and differential correlations

The task of our binary classifier is to assign one of two possible labels to a multi-dimensional pattern of activity. In most experimental situations, the two discrete labels are just particular instances of a continuous variable. For example, the direction of motion is a continuous variable, but the subject may be asked to discriminate between two particular directions (e.g., left vs right) in a motion discrimination task. Although we may measure neuronal activity only for particular values of a stimulus variable *s* (e.g. *s*_1_ and *s*_2_; *Δs* ≠ (*s*_1_ − *s*_2_)), the population tuning curve, **f**, is a continuously varying function of *s*, **f**(*s*). This mapping from the one-dimensional stimulus space 5 to the N-dimensional neuronal tuning space **f**(*s*) is assumed to be continuous and differentiable, and therefore 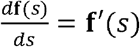 exists. We can assume that

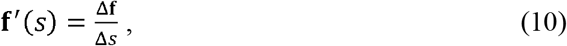

which is always true if *s*_1_ − *s*_2_ is small enough. In this case, the amount of encoded information (DP) can be obtained by substituting

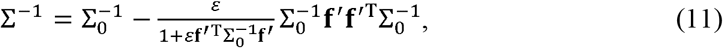

into 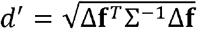

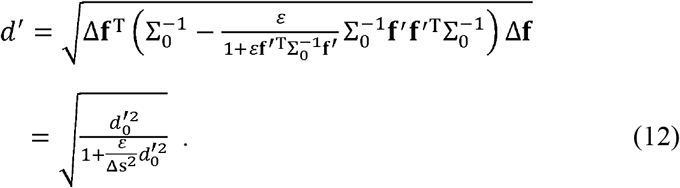

where 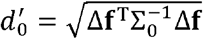.

Introducing Eq. (12) back into Eq. (3) gives us

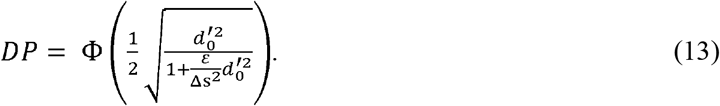

Therefore, when differential correlations are introduced into the system, the original 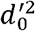 is reduced by a factor 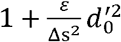. When the neuronal ensemble (N) is very large, this expression becomes

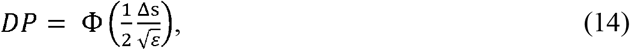

as 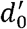 grows monotonically with N. In the presence of noise at the input stage, the decoding performance of a linear classifier will be constrained by the upper bound 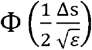, even when optimally reading-out the activity of many neurons simultaneously. The further the stimuli are apart, the higher the upper bound will be, and the larger the sensory noise is, the lower the upper bound becomes.

### 2. Comparison between theoretical and cross-validated decoding performance on experimental data

We evaluated how well decoding performance of the theoretical expression (DP_th_) agrees with that of the classifiers trained and tested on experimental data (DP_cv_). We plotted DPth against DP_c*v*_,and performed type-II regression (orthogonal linear regression) on the data. We assessed the similarity between the two measures of decoding performance by computing the percentage of the variance in DP_cv_ that can be explained by DP_th_ as *λ*_1_/(*λ*_1_ + *λ*_2_) 100, where *λ*_1_ and *λ*_2_ correspond to the variance captured by the first and second principal components of the DP_th_ vs DP_cv_ plot, respectively (Fig. S2). In order to account for possible idiosyncrasies from choosing a LDA when evaluating DP_cv_ and possible over-fitting on DP_cv_ due to the limited number of trials in our datasets, we also computed DP_cv_ using Logistic Regression (LR) on both the test and the training sets (Fig. S2). In addition to the percentage of explained variance as the goodness-of-fit metric for the linear fit (% explained variance or E.V.), we also considered the slope and intercept parameters as complementary goodness-of-fit metrics (Fig. S2) and obtained very similar overall results.

We compared how well the optimal expression (Eq. 4) approximated DP_cv_ with respect to the suboptimal expressions for DP_th_ (Eqs. 8 and 9) by plotting the ratio between the percentages of explained variance (Fig. S3A). We performed the same analysis after shuffling across trials the activity of each individual neuron for a given stimulus condition (Fig. S3B). Finally, we compared a linear discriminant analysis (LDA), a quadratic discriminant analysis (QDA) and a logistic regression (LR). Since LDA performed the best among the three (Fig. S4), we chose LDA for computing DP_cv_ in all analyses presented here. For a detailed description of how these fits were performed for each dataset, see section 5.

### 3. Conditioned bootstrapping method

We performed an analysis based on bootstrapping to test the effects of fluctuations in PS, PP and other statistical quantities of neuronal responses on the amount of encoded information (DP_cv_) and behavioral performance. The conditioned bootstrapping method involves two steps: (1) generating fluctuations in statistical quantities through bootstrapping, and (2) conditioning by selecting bootstrap samples that produce fluctuations in certain statistical features but not in the conditioned ones.

For a particular set of trials (sub-dataset, see below), bootstrap samples were generated by randomly selecting *M* trials (*M* being the total number of trials for this particular sub-dataset) with replacement. For each bootstrap sample we calculated the following quantities:

- Behavioral performance of the animal, denoted as B (see below for the definition of behavioral performance in each task).
- Theoretical decoding performance, DP_th_ (Eq. 4).
- Decoding performance of a cross-validated linear classifier (LDA), DP_cv_.
- Population Signal, PS, and projected precision, PP.
- Mean pairwise correlation, MPC, defined as the average of all pairwise correlations for a fixed stimulus condition.
- Global activity of the neuronal population, GA, defined as the mean neuronal activity across all neurons and trials.

We generated 10^4^ bootstrap samples (10^3^ when analyzing DP_cv_, as fitting the classifier is computationally expensive) and obtained a distribution of each quantity listed above. We will denote the obtained value of the statistical feature *i* for the bootstrap iteration *j* as *x_ij_*, and the fluctuation *x_ij_* of with respect to the median value of the distribution across bootstrap iterations as 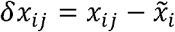, where 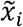 denotes the median of the distribution. We used the median rather than the mean so that the method works just as robustly for skewed distributions as for symmetrical distributions, although strongly skewed distributions were typically not observed (Fig. 2). It is important to note that bootstrap samples could include non-unique trials. When evaluating statistical features like the covariance matrix, if the number of unique trials is small, singular matrices could be generated. Nevertheless, the number of unique trials in all datasets is typically 10 times larger than the number of neurons used in the analysis (see next section). Therefore, the rank of the bootstrapped covariance matrix is always bound by the number of neurons rather than by the number of unique trials subsampled for each bootstrap iteration. In other words, the probability of generating covariance matrices of full rank (number of neurons) in 10^4^ bootstrap iterations is effectively one.

In order to assess what statistical features affect the amount of encoded information and behavioral performance the most, we evaluated the dependency of *ΔDP_cv_* and *ΔB* on *Δx_i_*, where, as above, *x_i_* represents a particular statistical feature of the neuronal responses. Here, dependency refers to various possible measurements, such as correlation. A dependency across bootstrap iterations between *ΔDP_cv_* and *Δx_i_* (referred here as *p*(*ΔDP_cv_, Δx_i_*)), and between *δB* and *Δx_i_* (*p*(*δB, δx_i_*)) could result from dependencies mediated by a third quantity *δx_k_*. For instance, Eq. (4) predicts that DP_cv_ depends exclusively on PS and PP, but since PP decreases as the ensemble’s noise (both individual and pairwise) increases, an inverse relationship between *Δ*MPC and *Δ*PP is expected. Therefore, we may find a correlation between *ΔDP_cv_* and *ΔMPC* as a result of their dependencies on *Δ*PP. To estimate the strengths of the dependencies *p*(*ΔDP_cv_, Δx_i_*) and *p*(*ΔB, Δx_i_*) that are not confounded by other variables, we developed a method for minimizing the correlation due to dependencies of *δDP_cv_* (or *δB*) and *δx_i_* on a third variable *δx_k_*. Among all generated bootstrap samples, we selected those that yielded *δx_k_* ≃ 0, i.e., *x_k_* values within the ± 10^th^ percentile of its median value. Based on the selected samples, we computed *p*(*ΔDP_cv, Δx_i__*|*Δx_k_* ≃ 0) (or *p*(*ΔB, Δx_i_*|*Δx_k_* ≃ 0)), i.e. the dependency between *ΔDP_cv_* (or *ΔB*) and *Δx_i_* conditioned on *δx_k_*. This method can easily be extended to multiple conditioning variables as long as there are enough numbers of bootstrap samples for the analysis that satisfy the conditions. Using this technique, we computed the following conditional dependencies:

- *p*(*δDP_cv_, δPS|δPP* ≃ 0, *δMPC* ≃ 0) and *p*(*δB,δPS*|*δPP* ≃ 0, *δMPC* ≃ 0)
- *p*(*δDP_cv_, δPP*|*δPP* ≃ 0, *δMPC* ≃ 0) and *p*(*δB, δPP*|*δPS* ≃ 0, *δMPC* ≃ 0)
- *p*(*δDP_cv_, δMPC*|*δPS* ≃ 0, *δPP* ≃ 0) and *p*(*δB, δMPC*|*δPS* ≃ 0, *δPP* ≃ 0)
- *p*(*δDP_cv_, δGA*|*δPS* ≃ 0, *δPP* ≃ 0) and *p*(*δB, δGA*|*δPS* ≃ 0, *δPP* ≃ 0)

To evaluate *p*(*δDP_cv_,δx_i_*|*δx_k_* ≃ 0, *δx_l_* ≃ 0) (Figs. 5, 8B and Fig. S8A) and *p*(*δB, δx_i_*|*δx_k_* ≃ 0, *δx_l_* ≃ 0) (Figs. 7, 8C and Figs. S6, S8B), we used 10^3^ and 10^4^ bootstrap iterations, respectively. With the conditioning criterion of ±10^th^ percentile around the median, about 4% of the total number of bootstrap iterations (40 for dependencies on DP_cv_ and 400 for B) satisfied the conditioning on both *x_k_* and *x_l_* from which to compute dependencies.

The dependencies *p*(*ΔDP_cv_, Δx_i_*|*Δx_k_* ≃ 0, *Δx_l_* ≃ 0) and *p*(*ΔB, Δx_i_*|*Δx_k_* ≃ 0, *Δx_l_* ≃ 0) were evaluated by using two metrics: (1) the Pearson correlation coefficient (Figs. S6B, S7G) and (2) the percent change (Figs. 5, 7, 8B,C, and Figs. S6A, S8). Percent change in *DP_cv_* with respect to *Δx_i_* (% change 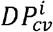) was defined as

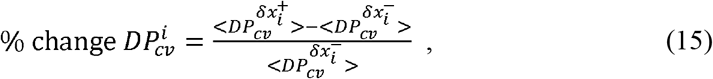

where 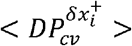 and 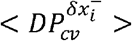 represent the mean DP_cv_ values for the bootstrap iterations that produced *Δx_i_* above 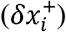 and below 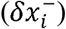 its median value 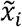, respectively. Similarly, percent change in *B* with respect to *δx_i_* (% change *B^i^*) was defined as

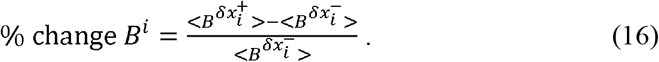

where 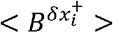 and 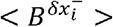 represent the mean *B* values for the bootstrap iterations that produced *Δx_i_* above 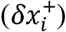 and below 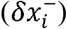 its median value 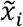, respectively. Because behavior (*B*) was quantified as the mean reaction time in the attentional task, the sign of *p*(*ΔB, Δx_i_*|*Δx_k_* ≃ 0, *Δx_l_* ≃ 0) was inverted so that negative fluctuations in *B* correspond to positive fluctuations in performance (Figs. 6 and 7B,C and Figs. S6A,B and S8B) for monkeys 2 and 3 (attentional task with recordings in LPFC 8a).

To confirm that our results still hold even if we impose a stricter conditioning criterion, we also computed % change *B* using bootstrap iterations that produced the conditioning variables *x_k_* and *x_l_* within ± 5^th^ percentile of their respective median values (100 bootstrap iterations fulfilled this conditioning criterion; see Fig. S6). We observed no substantial differences between the results from the two conditioning criteria.

The conditioning method employed here is an effective way of minimizing the portion of the dependency that is due to a third variable. One disadvantage of our approach is that, as the number of conditioning variables increases, the number of required bootstrap iterations increases exponentially. This is because, for a given iteration size, the number of bootstrap iterations that satisfy the simultaneous condition decreases. Because conditioning simultaneously on all features was computationally unfeasible due to the large number of bootstrap samples required, we also evaluated the conditioned dependencies described above but using GA instead of MPC as the conditioning variable when assessing the dependency between DP_cv_ and B on PS and PP. Specifically, we computed *p*(*δDP_cv_, δPS*|*δPP* ≃ 0, *ΔGA* ≃ 0), *p*(*ΔB, ΔPS*|*ΔPP* ≃ 0, *ΔGA* ≃ 0), *p*(*ΔDP_cv_, ΔPP*|*ΔPS* ≃ 0, *ΔGA* ≃ 0) and *p*(*ΔB, ΔPP*|*ΔPS* ≃ 0, *ΔGA* ≃ 0) and obtained qualitatively similar results to those depicted in Figs. 5, 7 (Supplementary Fig. S8).

For Fig. 3, the dependencies *p*(*ΔDP_cv_, ΔMPC*) and *p*(*ΔDP_cv_, ΔGA*) were evaluated using % change 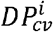 (Eq. 15) without conditioning on the rest of statistical features of the neuronal responses. For each monkey and ensemble size, the reported dependencies corresponded to the median value across independent sub-datasets (see detailed description of each experimental dataset in section 5). Statistical significance was calculated with a two-sided Wilcoxon signed-rank test, with which we tested whether the median of the distribution of independently obtained values was significantly greater or less than zero. Likewise, for Fig. 4, the dependencies *p*(*ΔPS, ΔMPC*), *p*(*ΔPS, ΔMPC*), *p*(*ΔPS, ΔMPC*) and *p*(*ΔPS, ΔMPC*) were evaluated with the Pearson correlation between the bootstrap fluctuations of these statistical features. For each ensemble size, the reported dependencies corresponded to the median values across independent sub-datasets and monkeys and statistical significance was calculated with a two-sided Wilcoxon signed-rank test as before. For Fig. 6 and S7G, the dependency *p*(*ΔB, ΔDP_cv_*) was evaluated by computing the Pearson correlation coefficient between *ΔDP_cv_* and *ΔB*. For each monkey and ensemble size, the median value across independent sub-datasets was reported and statistical significance was calculated with a two-sided Wilcoxon signed-rank test as for Figs. 3 and 4. Error bars correspond to the 25^th^ −75^th^ percentile of a distribution of medians obtained by the sampling with replacement from the distribution of independent values (bootstrap error bars; 1000 iterations).

### 4. Neural population model

We built a neural population model that captures the basic statistical properties of the neural responses that we observed experimentally. The model includes limited-range correlations, differential correlations, and realistic values for tuning sensitivity, firing rates and Fano factors. The model also generated choices based on optimal read out of information. We evaluated the simulation results by repeating all of the analyses described in the previous sections.

#### 4.1. Generative model

##### 4.1.1 Population activity model

Our neural population model consisted of *N* = 1000 neurons. Each neuron’s firing rate was modeled as a function of stimulus parameter *s* (such as motion direction) and stimulus strength, *v* (representing, for instance, motion coherence, which controls task difficulty). We considered two stimuli *s*_1_ = +1 and *s*_2_ = −1 (analogous to the two directions of motion around the discrimination boundary) at three stimulus strength levels, *v* = {0.16,0.32,0.48|. We defined mean firing rate *μ* of neuron *k* as a linear function of stimulus parameters (i.e., tuning curve) as

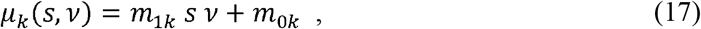

where *m*_1*k*_ and *m*_0*k*_ are the slope (sensitivity) and the baseline (spontaneous) firing rate of neuron *k*, respectively. The slope parameters *m*_1*k*_ were drawn randomly from a normal distribution centered at the origin with a standard deviation of *σ*_*m*_1__ = 1.3 and *m*_0*k*_ was drawn from a gamma distribution with shape and scale parameters set at 20 and 1, respectively. The parameters of the distributions were chosen to approximate empirical distributions. In what follows, we will use *μ_k_* (*s, v*) and *f_k_*(*s, v*) (defined earlier in Section 1) synonymously. We will use the terms spike count, firing rate and neuronal activity during a stimulus presentation interchangeably as the stimulus duration was set to 1 sec in our simulations.

Responses of neurons to identical stimuli vary from trial to trial, and the trial-by-trial variabilities are partially shared among neurons (noise correlation) (Cohen and Kohn 2011; Kohn et al. 2016). We incorporated noise correlations into our model in the form of limite-drange pairwise correlations between neurons *k* and *l* (Kanitscheider et al. 2015; Kohn et al. 2016) as

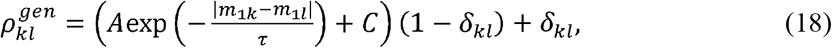

where *τ_kl_* is the Kronecker delta (*A* = 0.1, *C* = 0 and *τ* = 1 are used in our model). Then, we defined a generative covariance matrix as

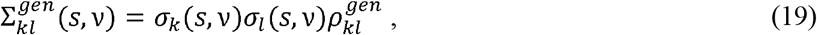

where 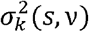 represents the trial-by-trial variance of neuron *k*. We used 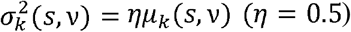 so that Fano Factors of the model neurons are within the rage of values typically found in experiments (Arandia-Romero et al. 2016; Nogueira et al. 2018; Shadlen and Newsome 1998). Note that 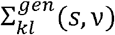 is not yet the full model’s covariance matrix, Σ_*kl*_(*s,v*), which includes differential correlations and other components as derived in the next section.

Based on ***μ***(*s,v*) and Σ^*gen*^, we generated neuronal activity as follows. First, we chose *s* and *v* pseudo-randomly for each trial of 1 sec duration such that there were 100 trials per stimulus condition (600 trials in total per simulated recording session). Then, we generated a preliminary response vector 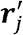 for each trial *j* as

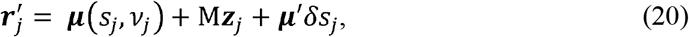

where 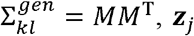, represents an *N*-dimensional vector whose elements are drawn from a zero-mean, unit-variance Gaussian distribution, ***μ***′ denotes the derivative of the population tuning curve (**f**), which equals ***m***_1_ in our model, and *δ_S_j__* indicates a common sensory noise term drawn from a Gaussian distribution with zero mean and variance 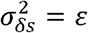. The last term ***μ***′*δ_S_j__* introduces differential correlations to the population activity (Kanitscheider et al. 2015; Moreno-Bote et al. 2014). The generated population activity pattern ***n**_j_* was calculated by applying multiplicative and additive global modulations to the vector 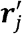 before putting it through a Poisson process that generated spike counts for each model neuron, i.e.

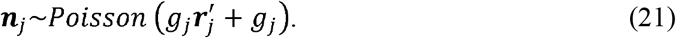

Here, *g_j_* is the global modulation factor drawn from a gamma distribution with scale and shape paramete *θ_g_* = 10^−5^ and *k_g_* = 10^5^ respectively (such that and *<g_i_* = 1 and 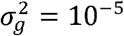). The global modulation incorporates the short and long timescale fluctuations in population activity often seen *in vivo* recordings (Arandia-Romero et al. 2016; Ecker et al. 2014; Goris, Movshon, and Simoncelli 2014; Lin et al. 2015). We generated a total of 10 simulated recording sessions that reproduced experimentally-realistic distributions of Fano factors and pairwise correlations (Figs. S7C,D).

##### 4.1.2 Behavioral decision model

We also modeled trial-by-trial behavioral choices using an optimal linear classifier that reads out the activity of the simulated population. For modelling behavior, the optimal classifier was derived from an analytical expression (Eqs. 22 and 23 below) rather than from data fitting since the simulated data had a limited number of trials (200 trials per stimulus intensity) and may result in overfitting. By introducing Eq. (17) into the analytical expression for the optimal linear classifier derived in Section 1, we obtain

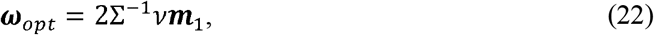

where Σ is the covariance matrix for our population model, which is given by

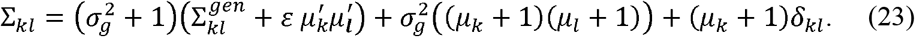

For each trial *j*, a decision variable was chosen as

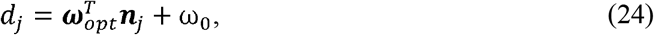

where 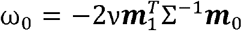. The behavioral choice *c* for trial *j* is, then, given by the sign of the decision variable *d*, i.e. *c_j_* = sign(*d_j_*). Behavioral performance was evaluated by the number of correct classifications over the total number of trials.

#### 4.2 Information encoded by the model

In this section, we derive a theoretical expression for decoding performance (DP_th_) of our model. In our model, 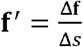, and therefore *d*′ is given by Eq. (12). When *N* → ∞

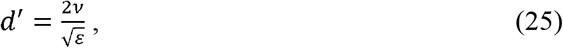

and therefore

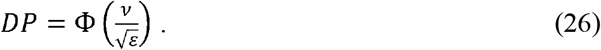

Equation (26) suggests that, when the input signal is noisy on a trial-by-trial basis, DP (the amount of information that can be extracted) of a very large neural ensemble does not saturate at 1.0. Instead, it will approach an asymptote determined by the signal to noise ratio of the input signal 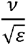 (see Figure S7F).

#### 4.3. Analysis of model simulation data

We performed the analyses described in Sections 2 and 3 on our simulated neuronal and behavioral data. First, we examined how well our theoretical decoding performance (DP_th_) approximated the cross-validated decoding performance (DP_cv_) of a linear classifier (LDA) evaluated on the simulated data, in the same way as we did for the *in vivo* recordings. For a particular ensemble size, we randomly selected a neuronal ensemble from a recording session and stimulus intensity and computed DP_th_ and DP_cv_. For each ensemble size, stimulus intensity and simulated recording session, we repeated this process 20 times. We then fitted a line to the relationship between DP_th_ and DP_cv_, and computed the percentage of variance in DP_cv_ that was explained by DP_th_, as described in Section 2 (Fig. S7E).

We also conducted the conditioned bootstrapping analyses on the simulated data using the method described in Section 3. We examined the effects of bootstrap fluctuations in statistical features of the population activity on the amount of encoded information and behavioral performance. For each bootstrap iteration, we selected an ensemble of a given size (2, 4, 6, 8 and 10 units) and calculated the following quantities: theoretical decoding performance (DP_th_), cross-validated decoding performance of a linear classifier (DP_cv_), population signal (PS), projected precision (PP), mean pairwise correlations (MPC), global activity (GA), and behavioral performance (B). Both DP_th_ and DP_cv_ were evaluated for inferring the stimulus (either *s*_1_ or *s*_2_) from the activity pattern of the population model. This process was repeated 20 times for each of the recording sessions and stimulus intensities. The dependencies of DP_cv_ and B on different features of the neural activity were quantified by % change in DP_cv_ and B as defined in Eqs. (15) and (16), as well as by Pearson correlation coefficients. The results were averaged over repetitions. For each ensemble size, we thus obtained 30 independent samples (10 surrogate recording sessions □ 3 stimulus intensities) and the median was reported. Statistical significance for the dependency between the variables *DP_cv_* (or *B*) and *δx_i_* through the calculation of *p*(*δDP_cv_, δx_i_|δx_k_* ≃ 0, *δx_l_* ≃ 0) (or *p*(*δB, δx_i_|δx_k_* ≃ 0, *δx_l_* ≃ 0)) for each ensemble size was calculated with a two-sided Wilcoxon signed-rank test, with which we tested whether the median of the distribution of 30 independently obtained values was significantly greater or less than zero. It is important to note that, when evaluating *p*(*δB, δx_i_|δx_k_* ≃ 0, *δx_l_* ≃ 0) as % change of surrogate behavior (Fig. 8C; Eq. (16)), small values can be obtained for relatively large surrogate ensemble sizes (N ~ 100) even when *p*(*δB,δx_i_|δx_k_* ≃ 0, *δx_l_* ≃ 0) is strong as evaluated by Pearson correlation. In our analysis, however, we prefer using % change of behavior because of its interpretability and robustness to outliers in the distribution of bootstrap fluctuations. The same bias could potentially arise when evaluating *p*(*δDP_cv_, δx_i_|δx_k_* ≃ 0, *δx_l_* ≃ 0) as % change of DP_cv_. Nevertheless, in this case, this effect is typically masked by the strong dependency between encoded information and PS/PP, a consequence of using the same neuronal population to compute all the relevant quantities.

### 5. Experimental Methods

Our theory was tested on three different datasets obtained from four monkeys, each of which involved simultaneously recorded units (from 2 to ~50 units). One dataset involved pairs of middle temporal (MT) neurons from a monkey (monkey 1) performing a coarse direction discrimination task (Zohary et al. 1994). The second dataset involved lateral prefrontal cortex (LPFC, area 8a) neurons recorded from two monkeys (monkeys 2 and 3) that were trained to perform an attentional task (Tremblay et al. 2015). The final dataset involved recordings from groups of MT neurons in a monkey (monkey 4) trained to perform a fine direction discrimination task. The details for each experiment are described below.

#### 5.1 Coarse direction discrimination task with recordings in area MT

This dataset has been previously described (Zohary et al. 1994), and is freely available at the Neural Signal Archive (http://www.neuralsignal.org, nsa2004.2)). Here we provide a brief summary.

##### 5.1.1. Subjects and recordings

Data from one adult macaque monkey (*Macacca mulatta*) are included in this dataset. The animal was trained to perform a coarse direction discrimination task (Britten et al. 1992), as described further below. Pairs of single neurons were recorded in area MT during this experiment. The animals were maintained in accordance with guidelines set by the U.S. Department of Health and Human Services (NIH) Guide for the Care and Use of Laboratory Animals. Electrophysiological recordings were made with tungsten microelectrodes (Micro Probe, Potomac, MD). Action potentials from single neurons were discriminated using an online spike sorting system. In total, 82 well-isolated neurons were recorded (41 pairs of single units).

##### 5.1.2 Experimental task

The monkey performed a coarse direction discrimination task, in which a noisy random-dot motion stimulus moved in one of two opposite directions. In each trial, a stimulus was presented for 2 seconds and covered the receptive fields of both neurons in a pair. Two motion directions were defined: preferred and null. the presented direction of motion was either the preferred or null direction of the recorded neurons. If the preferred directions of the two neurons differed substantially, the axis of discrimination was set to the preferred-null axis of the best-responding. The null direction was defined as the direction opposite to the preferred direction. The strength of the motion signal was manipulated by controlling the fraction of dots that moved coherently from one frame to the next (motion coherence), while the remaining ‘noise’ dots were randomly replotted on each video update. When motion coherence was 0%, all dots moved randomly, and there was no coherent direction in the net motion. When motion coherence was 100%, all dots moved coherently in either the preferred or null direction for the pair of neurons. A range of coherences between 0 and 100% was used to adjust task difficulty and measure neural and behavioral sensitivity. In each trial, the monkey was presented randomly with either the preferred or null direction of motion at a particular coherence. The monkey’s task was to report the direction of the net motion by making a saccade to one of two choice targets. The psychometric function averaged across sessions is shown in Fig. S6A.

Each trial started with the appearance of a fixation point. After the monkey held its gaze on the fixation target for 300 ms, the random-dot motion stimulus was presented. The animal was required to maintain fixation during stimulus presentation so that stimulus was presented over the receptive fields of the recorded neurons. After 2 seconds of stimulus presentation, the random dot pattern and the fixation point disappeared, and two choice targets appeared on the screen, corresponding to the two possible directions of motions (preferred or null) (see Figure 1A). The monkey made a saccadic eye movement to one of the targets to report its perceived direction of motion, and was provided a fluid reward for the correct choice, except for 0% coherence trials for which the monkey was rewarded randomly on 50% of trials.

##### 5.1.3 Neuronal data analysis

Data from this experiment (Zohary et al. 1994) consisted of 41 recording sessions in which pairs of single-units were simultaneously recorded in area MT. Note that only sessions with random-dot motion stimuli generated from a variable seed were used.

Since stimulus strength was controlled by the motion coherence parameter, we subdivided each recording session according to the coherence presented in each trial. Trials belonging to the 0% coherence condition were discarded because stimulus identity and correct behavioral performance were not defined for this condition. In order to evaluate how accurate our analytical approximation was for the amount of information encoded by the neural population, we plotted DP_th_ against DP_cv_ for each recorded pair of neurons and coherence condition and fitted the data points with a type-II linear regression. The percentage of variance explained by the first principal component of the data represented the goodness-of-fit (see section 2).

For the analysis involving behavior, we also discarded trials with coherences that elicited a mean behavioral performance greater than 98% correct to avoid ceiling effects (25.6%, 51.2% and 99.9% coherence were eliminated by this criterion). This was done because the bootstrap distributions for behavioral performance (B) would have very low variability for these high coherences, and therefore calculating *p*(*δB, δx_i_|δx_k_* ≃ 0, *δx_l_* ≃ 0) may be highly biased or even undefined. The criterion for discarding easy trials (98% percent correct) corresponds to eliminating conditions that are more than two standard deviations away from the mean of a cumulative Gaussian fit, since Φ(2*σ*) ≃ 0.98. After splitting recording sessions by coherence and discarding 0% and high coherences as described above, we obtained 187 independent sub-datasets, each of which corresponded to a set of trials for a particular coherence level and recording session. The mean number of trials per sub-dataset was 75, ranging from 30 to 231 trials. For the analysis, we used a trial-by-trial population activity vector whose entries corresponded to the spike count from the entire 2 second stimulus duration for each neuron.

From each sub-dataset, we generated bootstrap distributions for the following quantities: theoretical decoding performance (DP_th_), cross-validated (trained and tested on different sets of trials) decoding performance of a linear classifier (DP_cv_), population signal (PS), projected precision (PP), mean pairwise correlation (MPC), global activity (GA), and behavioral performance (B). Behavioral performance was defined as the fraction of correct choices of the monkey on each sub-dataset. Both DP_th_ and DP_cv_ were calculated by inferring the motion direction (preferred or null) presented to the monkey on a trial-by-trial basis from the simultaneously recorded activity of a neuronal pair.

Significance for the dependency *p*(*δDP_cv_, δx_i_|δx_k_* ≃ 0, *δx_l_* ≃ 0) (or *p*(*δB,δx_i_|δx_k_* ≃ 0, *δx_l_* ≃ 0)) for the whole experiment (187 independent sub-datasets) was calculated with a two-sided Wilcoxon signed-rank test, with which we tested whether the median of the distribution of 187 independently obtained values was significantly greater or less than zero.

#### 5.2. Attentional task with recordings in LPFC (area 8a)

This dataset is described in detail in (Tremblay et al. 2015). Here we provide a brief summary.

##### 5.2.1. Subjects and recordings

Two male monkeys (*Macaca fascicularis*), both 6 years old (Monkey “F”, 5.8 Kg; Monkey “JL”, 7.5 Kg), contributed to this dataset. In each monkey, a 96-channel “Utah” multielectrode array (Blackrock Microsystems, Utah, USA) was chronically implanted in the left caudal lateral prefrontal cortex. The multielectrode array was inserted on the prearcuate convexity posterior to the caudal end of the principal sulcus and anterior to the arcuate sulcus, a region cytoarchitectonically known as area 8a. The extracted spikes and associated waveforms were sorted offline using both manual and semi-automatic techniques (Offline sorter, Plexon Inc., TX, USA). All procedures were in accordance with the Canadian Council of Animal Care guidelines and were preapproved by the McGill University Animal Care Committee. Neither animal was sacrificed for the purpose of this study.

##### 5.2.2. Experimental task

The monkeys were trained to covertly sustain attention to one of four Gabor stimuli (target) presented on a screen while ignoring the other three Gabor stimuli (distractors) (Figure 1B). At the beginning of each trial, a cue indicated which of the four Gabor stimuli was the target (cue period, 363 ms). After the cue period, all four Gabor stimuli appeared on the screen, which marked the start of the attentional period. The attentional period ended after a variable delay (585-1755 ms) when one or two Gabor stimuli changed orientation by 90°. Three different trial types were randomly interleaved. In “Target” trials, the target Gabor changed orientation, prompting the monkey to make a saccade towards the target within 400 milliseconds to get a reward (fruit juice). In “Distractor” trials, the orientation change occurred in the Gabor location diagonally opposite to the cued location. Monkeys had to withhold saccades to the distractor and maintain fixation for the trial to be correct. In “Target + Distractor” trials, the orientation of the target and the distractor diagonally opposite to the target changed simultaneously. In this case, the monkey had to make a saccade towards the target, not to the distractor, to get a reward. On average, monkeys completed ~1000 trials per session (Fig S5B,C). Only correct “Target” trials were used in the analysis reported here. The other trial types were necessary for keeping the performance of the animals unbiased.

##### 5.2.3. Neuronal data analysis

The mean number of correct “Target” trials per recording session was 207 (range: 172 to 224) for monkey “JL” and 221 trials (range: 198 to 246) for monkey “F”. The mean number of simultaneously recorded units was 56 (range: 53 to 61) for monkey “JL” and 54 (range: 44 to 66) for monkey “F”. Neuronal recordings included both single and multiunits. The mean percentage of single units over the total number of simultaneously recorded units was 44% and 40% for monkeys “JL” and “F”, respectively. Because units with very low firing rates precluded reliable statistical analysis, we excluded units with firing rates below 1 Hz for all subsequent analyses. After this exclusion, the mean number of simultaneously recorded units was 51 (range: 50 to 53) for monkey “JL” and 50 (range: 42 to 59) for monkey “F”. The shortest attentional period used was 585 ms; therefore, we defined a fixed attentional time window of 585 ms starting at the end of the cue period. In this way, the firing rate of all units was calculated over the same time window.

To maximize the statistical power of our analysis, we created a larger number of independent sub-datasets as follows. For each recording session, we randomly selected 21, 10, 7, 5, and 4 non-overlapping ensembles of size 2, 4, 6, 8, and 10, respectively, and repeated this process 5 times. Since the smallest number of simultaneously recorded units for both monkeys was 42 units, we chose values that maximized the number of non-overlapping ensembles for each size.

In order to evaluate how accurate our analytical approximation was for the amount of information encoded by the neural population for a particular ensemble size, we plotted DP_th_ against DP_cv_ for each sub-dataset and fitted the data points with a type-II linear regression. The percentage of variance explained by the first principal component of the data represented the goodness-of-fit of our approximation (see section 2).

For each sub-dataset and bootstrap iteration, we calculated the following quantities: theoretical decoding performance (DP_th_), cross-validated decoding performance of a linear classifier (DP_cv_), population signal (PS), projected precision (PP), mean pairwise correlation (MPC), global activity (GA), and behavioral performance (B). Behavioral performance (B) was quantified as the mean reaction time (RT) across trials (either the original sub-dataset or a bootstrap iteration) (see Figure S5B,C for RT distributions from monkeys 2 and 3, respectively). Both DP_th_ and DP_cv_ were calculated by inferring the monkey’s location of attention on a trial-by-trial basis from the simultaneously recorded activity of an ensemble. For decoding purposes, we considered two binary classification tasks. In the first task, the decoder classified the locus of attention as being on the right or the left side of the screen. In the second task, the decoder classified the locus of attention as being on the upper half or the lower half of the screen. The reported DP_cv_ (and DP_th_) values are averages over the two decoding tasks.

For a given sub-dataset, the reported dependency relationship *p*(*δDP_cv_,δx_i_|δx_k_* ≃0, *δ_l_* ≃ 0) (or *p*(*δB,δx_i_|δx_k_* ≃0, *δ_l_* ≃ 0)) was evaluated as the mean across the 5 iterations and the two classification tasks (vertical and horizontal). For each ensemble size and monkey, the reported dependency *p*(*δDP_cv_,δx_i_|δx_k_* ≃0, *δ_l_* ≃ 0(or *p*(*δB,δx_i_|δx_k_* ≃0, *δ_l_* ≃ 0)) was the median across recording sessions and non-overlapping ensembles of units (independent sub-datasets). We obtained 84, 40, 28, 20 and 16 (the number of non-overlapping ensembles □ 4 recording sessions per monkey) independent values for ensemble sizes of 2, 4, 6, 8, and 10 units, respectively, and assessed the median and tested significance. Significance was calculated with a two-sided Wilcoxon signed-rank test of whether the median of the distribution of independently obtained values for each ensemble size and monkey was significantly above or below zero.

#### 5.3. Fine direction discrimination task with recordings in MT

This dataset was obtained specifically for this analysis, as part of a larger series of ongoing studies of how MT neurons represent local motion in the presence of background optic flow.

##### 5.3.1. Subjects and recordings

One adult macaque monkey *(Macaca mulatta)* was used in this experiment. The animal was surgically implanted with a circular head holding device, a scleral coil for measuring eye movements, and a recording grid (Gu et al. 2006; Gu, Angelaki, and DeAngelis 2008). The animal was trained to perform a fine direction discrimination task (described below) with water as a reward for correct performance. Eye movements were measured and controlled at all times. Neuronal activity in MT/V5 was recorded with 24-channel linear electrode arrays (V-probes, Plexon Inc). Spike waveforms were acquired by a Blackrock Cerebus system. All experimental procedures conformed to National Institutes of Health guidelines and were approved by the University Committee on Animal Resources at the University of Rochester.

In each recording session, a V-probe was inserted into MT/V5 and was allowed to settle for ~30 minutes. We then performed standard tests (DeAngelis and Uka 2003) to map receptive fields and to measure the direction tuning of the recorded units. These measurements were used to determine the location and the size of the stimulus aperture such that it was larger than the receptive fields of the units under study. Note, however, that stimulus motion was not tailored to the preferred directions and speeds of the recorded neurons; a fixed set of stimulus velocities was used across sessions for the fine discrimination task. A total of 75 units were recorded across 3 sessions. The mean number of simultaneously recorded units was 25, ranging from 24 to 27. Most recordings included multi-unit activity as well as 2-4 well-isolated single units. Because units with very low firing rates precluded reliable statistical analysis, we excluded units with firing rates below 1 Hz for all subsequent analyses. Despite the exclusion, the mean number of simultaneously recorded units remained at 25 (ranging from 24 to 27).

##### 5.3.2. Experimental task

The monkey was trained to perform a fine direction discrimination task. A pattern of random dots, presented within a circular aperture, moved upward in the visual field with either a rightward or leftward component. The animal’s task was to report whether the perceived motion was up-right or up-left by making a saccade to one of two choice targets. The motion stimulus was presented at 100% coherence in one visual hemi-field, and was localized and sized according to the receptive fields of the recorded units. Stimulus motion within the aperture followed a Gaussian velocity profile with a standard deviation of 333 ms and a duration of 2 s. Once the monkey fixated for 200 ms, the motion stimulus appeared and began to move. The stimulus was presented stereoscopically at zero disparity, such that motion appeared in the plane of the display. The experimental protocol involved 7 directions of motion relative to vertical: −12°, −6°, −3°, 0°, 3°, 6° and 12°, where negative and positive values correspond to leftward and rightward motion, respectively. After stimulus presentation was completed, two saccade targets appeared, 5 deg. to the right and left of the fixation target, and the monkey reported his perceived direction of motion by making a saccade to one of the targets (Fig. 1C). Because stimuli were presented at 100% coherence, task difficulty was controlled by the direction of motion with respect to vertical (see Figure S6D). The mean number of trials per session was 742 (range: 735 to 756). In each recording session, the same number of trials were presented for each direction of motion.

##### 5.3.3. Neuronal data analysis

Since direction of motion controlled the difficulty of the task, this parameter was considered to be the stimulus strength. To control for stimulus strength, we divided data from each recording session according to motion direction. The task of our decoder is to correctly classify each trial as having rightward or leftward motion direction; therefore, trials from each recording session were split into 4 independent sub-datasets (±12°, ±6°, ±3° and 0° stimulus directions relative to vertical). As described above for the coarse discrimination task, we subsequently discarded stimulus conditions that were either ambiguous (0° motion direction; correct behavioral performance is not defined) or too easy (±12° directions; mean behavioral performance ≥ 0.98). The analysis was thus performed on 6 independent sub-datasets (3 recording sessions □ 2 absolute motion directions). To quantify neuronal activity, we computed firing rates in a time window that included ±1 standard deviation around the peak of the Gaussian velocity profile of the stimulus (666 ms window width).

To increase statistical power, we created a larger number of independent sub-datasets by sampling randomly non-overlapping ensembles of particular sizes. Namely, we considered ensemble sizes of 2, 4, 6, 8, and 10, which yielded 12, 6, 4, 3, and 2 non-overlapping ensembles of units, respectively. We repeated this sampling procedure 20 times. Since the smallest number of simultaneously recorded units was 24, we chose values that maximized the number of non-overlapping ensembles for each size.

We evaluated how accurate DP_th_ was with respect to DP_cv_ by following the same procedure as for monkeys 2 and 3 (see previous section). For each sub-dataset and bootstrap iteration, we calculated the following quantities: theoretical decoding performance (DP_th_), cross-validated decoding performance of a trained and tested linear classifier (DP_cv_), population signal (PS), projected precision (PP), mean pairwise correlation (MPC), global activity (GA), and behavioral performance (B). For this task, behavioral performance was defined as the fraction of correct choices for that particular sub-dataset. Both DP_th_ and DP_cv_ were calculated by inferring whether motion direction was leftward or rightward of vertical for each trial based on the neuronal activity pattern. For a particular randomly constructed ensemble, the reported dependency relationship *p*(*δDP_cv_,δx_i_|δx_k_* ≃0, *δ_l_* ≃ 0) (or *p*(*δB,δx_i_|δx_k_* ≃ 0,*δx_l_* ≃ 0)) was evaluated as the mean across the 20 iterations. For each ensemble size, the reported dependency *p*(*δDP_cv_,δx_i_|δx_k_* ≃0, *δ_l_* ≃ 0) (or *p*(*δB,δx_i_|δx_k_* ≃ 0,*δx_l_* ≃ 0)) was the median across recording sessions, stimulus strengths, and non-overlapping ensembles of units. We used 72, 36, 24, 18 and 12 (the number of non-overlapping ensembles □ 3 independent recording sessions □ 2 stimulus strengths) independent values to assess the median and test its significance for ensemble sizes of 2, 4, 6, 8, and 10 units, respectively. Significance was calculated with a two-sided Wilcoxon signed-rank test for whether the median of the distribution of independent values for each ensemble size was significantly greater or less than zero.

## Quantification and statistical analysis

See previous section.

## Data and software availability

The datasets generated in this study and the code used for their analysis are available from the corresponding author upon reasonable request.

