## Supplemental Information for "Identifying the most influential features of neural population responses for information encoding and behavior"

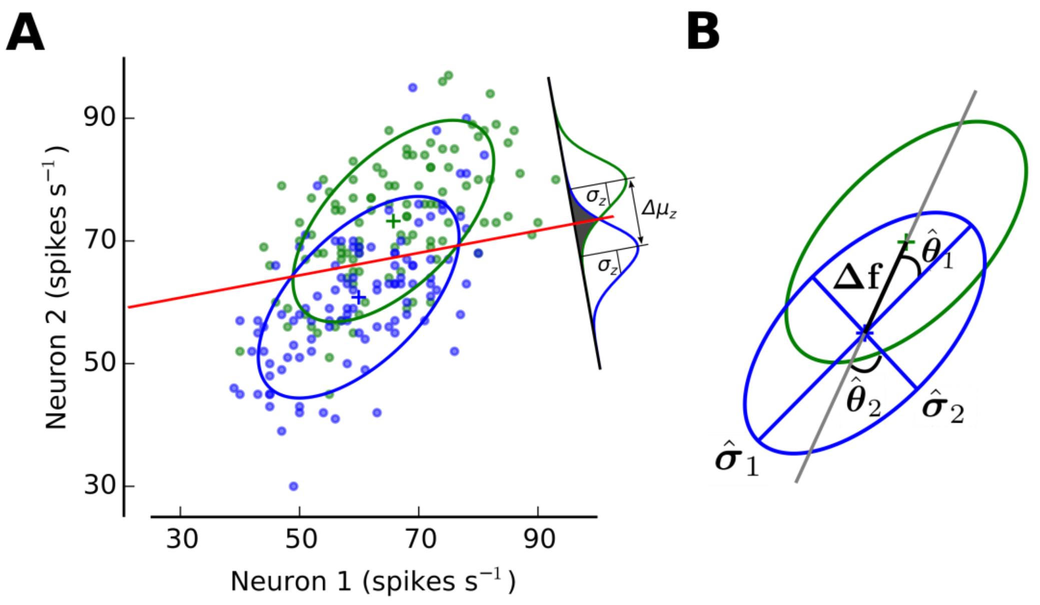


**Figure S1. The information encoded by a neuronal ensemble is fully characterized by population signal and projected precision. Related to STAR Methods.**

**(A)** The trial-by-trial joint activity of a network consisting of *N* neurons can be characterized by an ellipsoid embedded in an *N*-dimensional space (a representative population of two MT neurons from monkey 4 is shown here; see STAR Methods). The covariance matrix and mean activity of the population determine the shape and location of the ellipsoids corresponding to stimulus 1 ($s_1$; green) and stimulus 2 ($s_2$; blue). The linear classifier is characterized by a linear boundary that separates the two clouds of points as well as possible (red line). The difference in mean value for the decision variable $z$ and its standard deviation are represented by $\Delta\mu_z$ and $\sigma_z$, respectively (signal-to-noise ratio of d′). **(B)** Graphical depiction of population signal (PS) and projected precision (PP). PS is defined as the norm of the tuning vector, $\mathbf{\Delta f}$, which corresponds to the distance between the mean responses associated with $s_1$ and with $s_2$ (distance between the centers of the green and blue ellipsoids). PP is calculated from the angles ($\hat{\theta}_i$) between each eigenvector of the covariance matrix and the tuning vector ($\mathbf{\Delta f}$) (orientation of each axis of the ellipsoid with respect to $\mathbf{\Delta f}$) as well as from their eigenvalues ($\hat{\sigma}_i^2$; length of each axis). Greater information is encoded when the longest axis of the ellipsoid becomes orthogonal to the direction of $\mathbf{\Delta f}$.

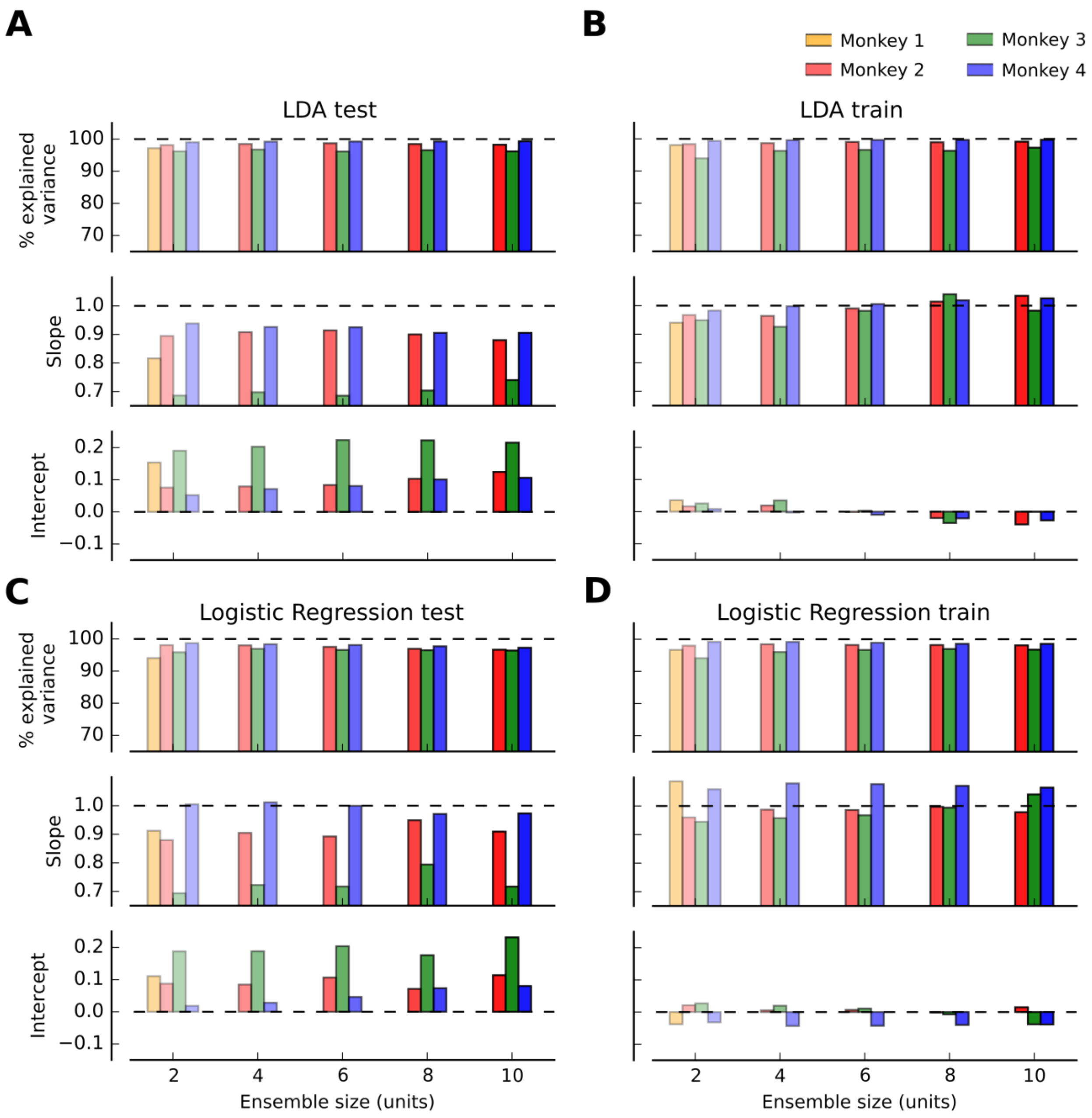


**Figure S2. The theoretical expression provides excellent fits for different linear classifiers and goodness-of-fit metrics. Related to STAR Methods.**

**(A)** The slope and intercept parameters of the linear fit (Type II regression, see STAR Methods) between the theoretical and the cross-validated decoding performance ($DP_{th}$ and $DP_{cv}$) are relatively close to 1.0 and 0.0, respectively, for a Linear Discriminant Analysis (LDA) on all ensemble sizes and monkeys. Deviations from 1.0 (slope) and 0.0 (intercept) can be explained by the limited number of trials used to train and test the LDA, which produces values of DP below

0.5 in some cases. (**B**) All goodness-of-fit metrics (% explained variance, slope and intercept) are improved when the training set is used to test the performance of LDA. (**C**) When using Logistic Regression (LR) instead of LDA, the fitting results are similar to panel (A). (**D**) As in panel (B), when using the training set to evaluate the performance of LR, all goodness-of-fit metrics are improved with respect to panel (C).

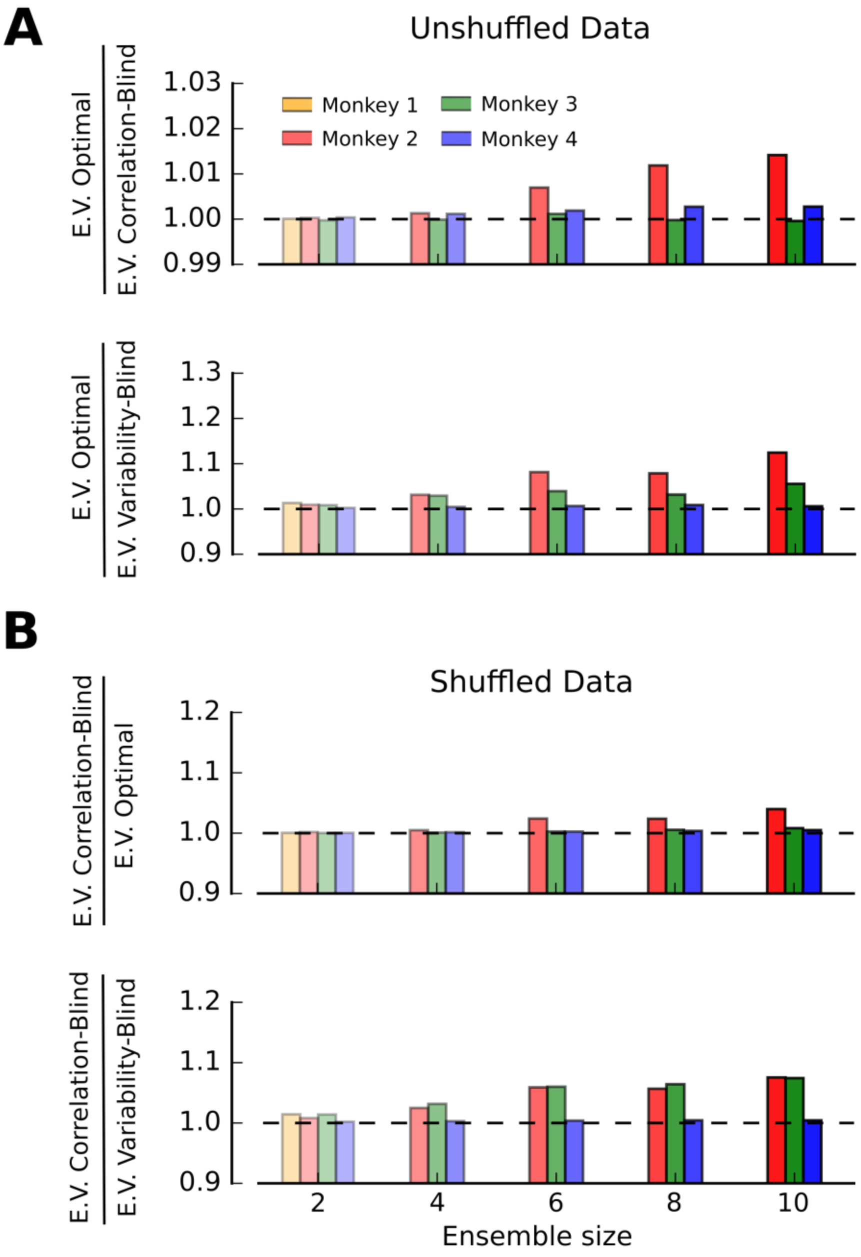


**Figure S3. The expression for the optimal classifier is the best approximation to the amount of encoded information. Related to STAR Methods.**

**(A)** Ratio between percentage of explained variance (E.V.) of the optimal classifier and the correlation-blind classifier (top panel) and between the optimal and the variability-blind classifier (bottom panel; see STAR Methods). The percentage of explained variance determines how good a given analytical expression of $DP_{th}$ approximates the performance of a cross-validated linear classifier $DP_{cv}$ (see STAR Methods). For all monkeys (mean across all ensemble sizes: 2, 4, 6, 8 and 10 units) the theoretical expression for the optimal classifier (Eq. 1) is the best approximation for the amount of information encoded by the neural population as expressed by $DP_{cv}$. (**B**) Ratio between percentage of explained variance (E.V.) of the correlation-blind classifier and the optimal classifier (top panel) and between the correlation-blind and the variability-blind classifier (bottom panel) after shuffling the activity of each neuron across trials for a fixed condition. The amount of encoded information when pairwise correlations are removed is better approximated by the correlation-blind classifier than by the optimal classifier (before shuffling) and the variability-blind classifier for all monkeys.

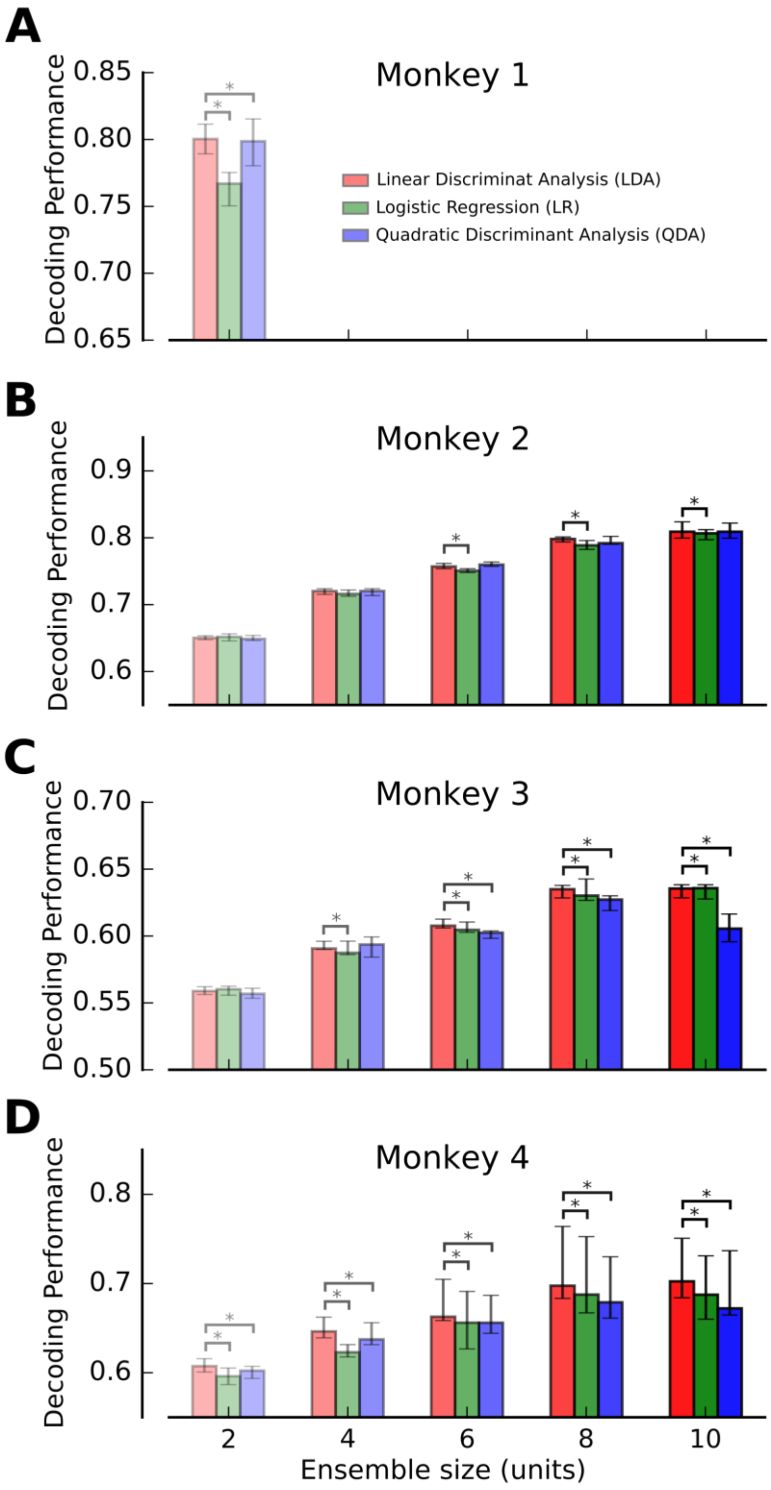


**Figure S4. Linear discriminant analysis outperforms logistic regression and quadratic discriminant analysis. Related to STAR Methods.**

**(A)** Median decoding performance (DP) for linear discriminant analysis (LDA), logistic regression (LR), and quadratic discriminant analysis (QDA) (5-fold cross-validation) for an ensemble size of 2 units from monkey 1. LDA produces a larger DP than both LR ($P$ = $6.2 \times 10^{-13}$, Wilcoxon signed rank test) and QDA ($P$ = $2.4 \times 10^{-3}$). Note that significance of the differences is strong due to pairing of the data despite the relatively large error bars displayed. (**B, C**) Equivalent to (A) for monkeys 2 and 3 and for a range of ensemble sizes (2, 4, 6, 8 and 10 units). As in (A), LDA produces the largest mean DP of the three decoders. (**D**) Analogous results for monkey 4. In all panels error bars denote $25^{th}$ -$75^{th}$ percentile of the distribution of bootstrap medians and * indicates significance in the reported differences ($P < 0.05$).

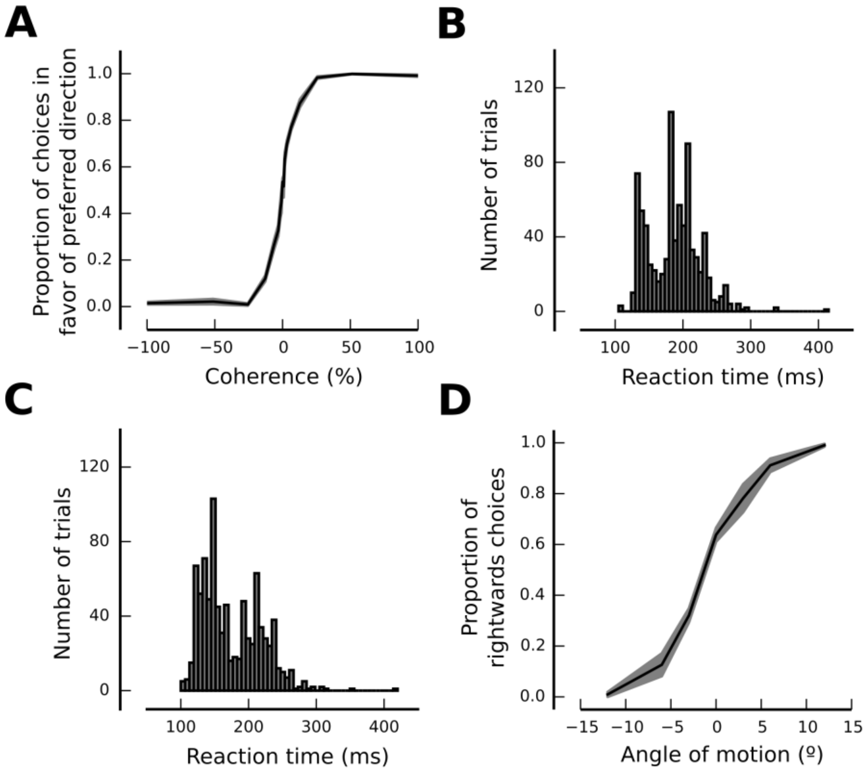


**Figure S5. Definitions of behavioral performance for each task. Related to STAR Methods.**

**(A)** Psychometric curve averaged across recording sessions for monkey 1. Proportion of choices in favor of the preferred direction as a function of motion coherence (positive values of coherence correspond to the preferred direction of motion). Grey region: s.e.m. **(B, C)** Distributions of reaction times for monkeys 2 and 3. For the attentional task, reaction time is defined as mean time (across trials in a particular dataset) from the change in stimulus orientation until the saccade to the cued Gabor pattern. (**D**) Psychometric curve averaged across recording sessions for monkey 4. Proportion of rightward choices as a function of the direction of motion with respect to vertical (positive and negative values denote motions with a rightward and leftward component, respectively). Grey region: s.e.m.

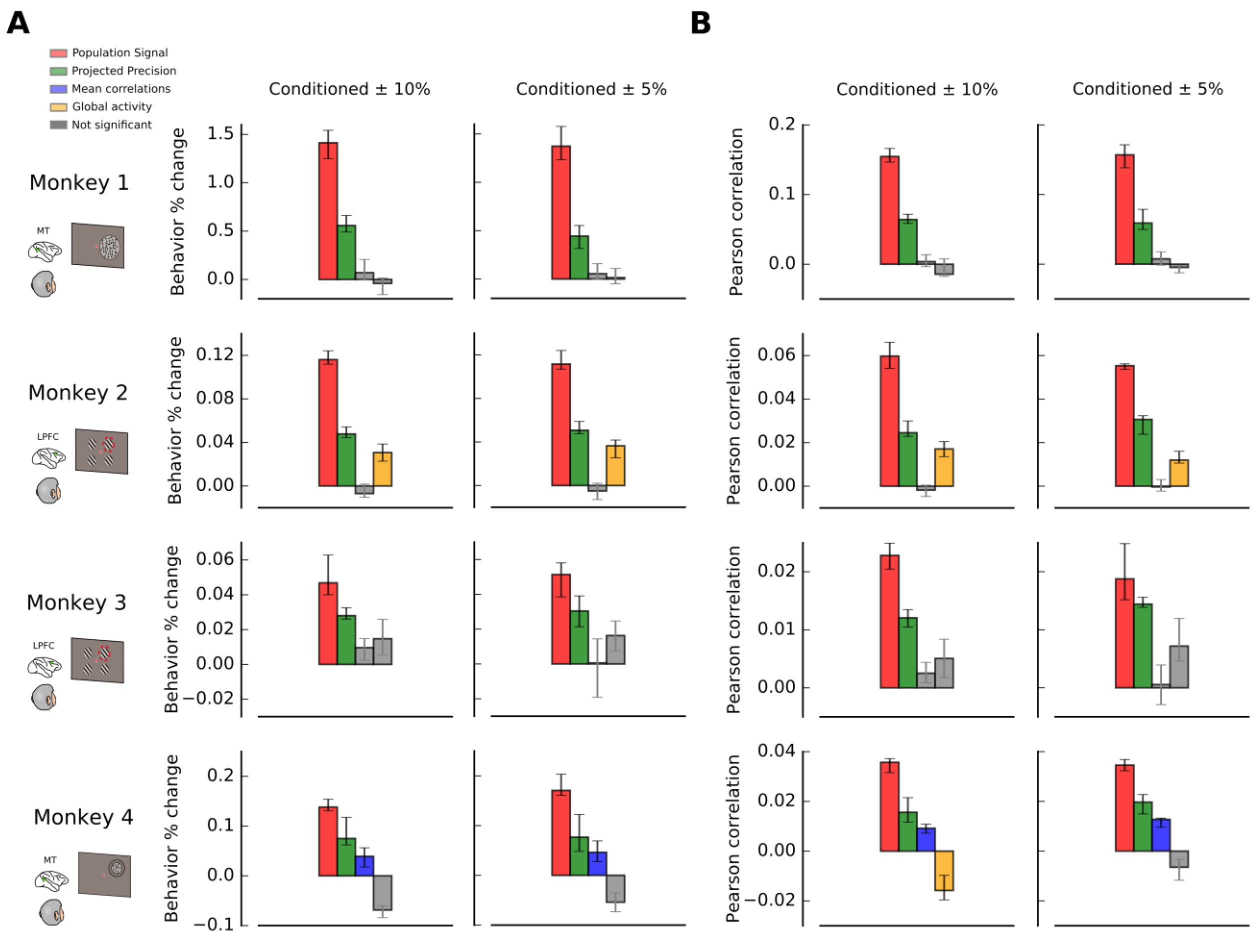


**Figure S6. Results shown in Fig. 7 are robust to different conditioning parameters and metrics. Related to Fig. 7.**

**(A)** Equivalent to Fig. 7, but averaged across ensemble sizes. Percentage change in behavior (see STAR Methods) is plotted for bootstrap fluctuations of each statistical feature (conditioned bootstrapping method; see STAR Methods), and results are shown for different conditioning criteria: ± 10% (left column) and ± 5% (right column) from the median value of the bootstrap distribution. (**B**) Same as (A) but using Pearson correlation between bootstrap fluctuations of the different features of neural activity and behavioral performance instead of percentage change. Data are shown for the same two conditioning criteria and averaged across ensemble sizes. Error bars correspond to $25^{th}$ -$75^{th}$ percentile of the distribution of bootstrap medians and significant deviations from zero (colored bars) are calculated by a Wilcoxon signed rank test (not significant if $P > 0.05$).

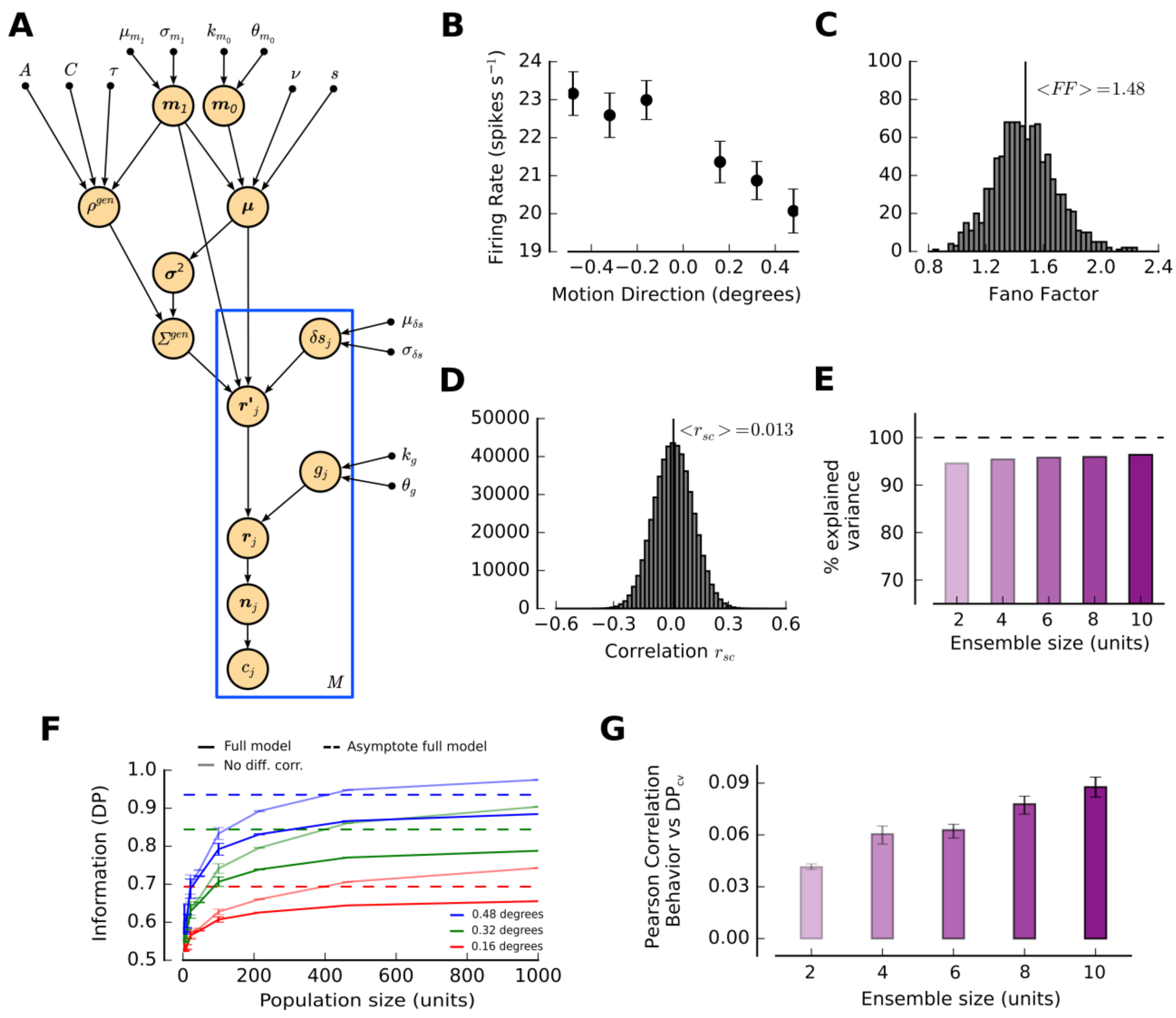


**Figure S7. An experimentally-constrained population model replicates experimental results. Related to Fig. 8.**

(**A**) Graphical model depicting, in full detail, the generative process underlying the surrogate datasets (see STAR Methods). (**B**) Tuning curve of an example neuron used in the model. This neuron's firing rate signals the direction of motion with a linear tuning curve that is corrupted by noise. (**C**) Distribution of Fano Factors for all units in an example surrogate 'session' of the model. The mean value of the distribution is in rough agreement with empirical values of Fano factors found in *in vivo* recordings. (**D**) Distribution of pairwise correlations of the spike counts of all neuronal pairs in an example surrogate 'session' of the model. The mean value of the distribution is compatible with the empirical values of mean pairwise correlations found in *in vivo* recordings. (**E**) The theoretical decoding performance of a linear classifier ($DP_{th}$) was a very good approximation of the cross-validated decoding performance ($DP_{cv}$) on the model data (see STAR Methods and Fig. S2). (**F**) Amount of encoded information ($DP_{cv}$) as a function of the network size for the three different stimulus intensities in the model. When differential correlations are not included in the model (light colored lines), information grows without limit for all of the stimulus values ($DP_{cv}$ = 1.0 upper limit for information in a classification task). When differential correlations are included in the model (darker colored lines), $DP_{cv}$ is lower and reaches an asymptote (dashed colored lines) for large network sizes. (**G**) Pearson correlation between bootstrap fluctuations in the amount of encoded information ($DP_{cv}$) and behavioral performance of the virtual agent. The model accounts for the weak coupling between these two quantities (see Fig. 6).

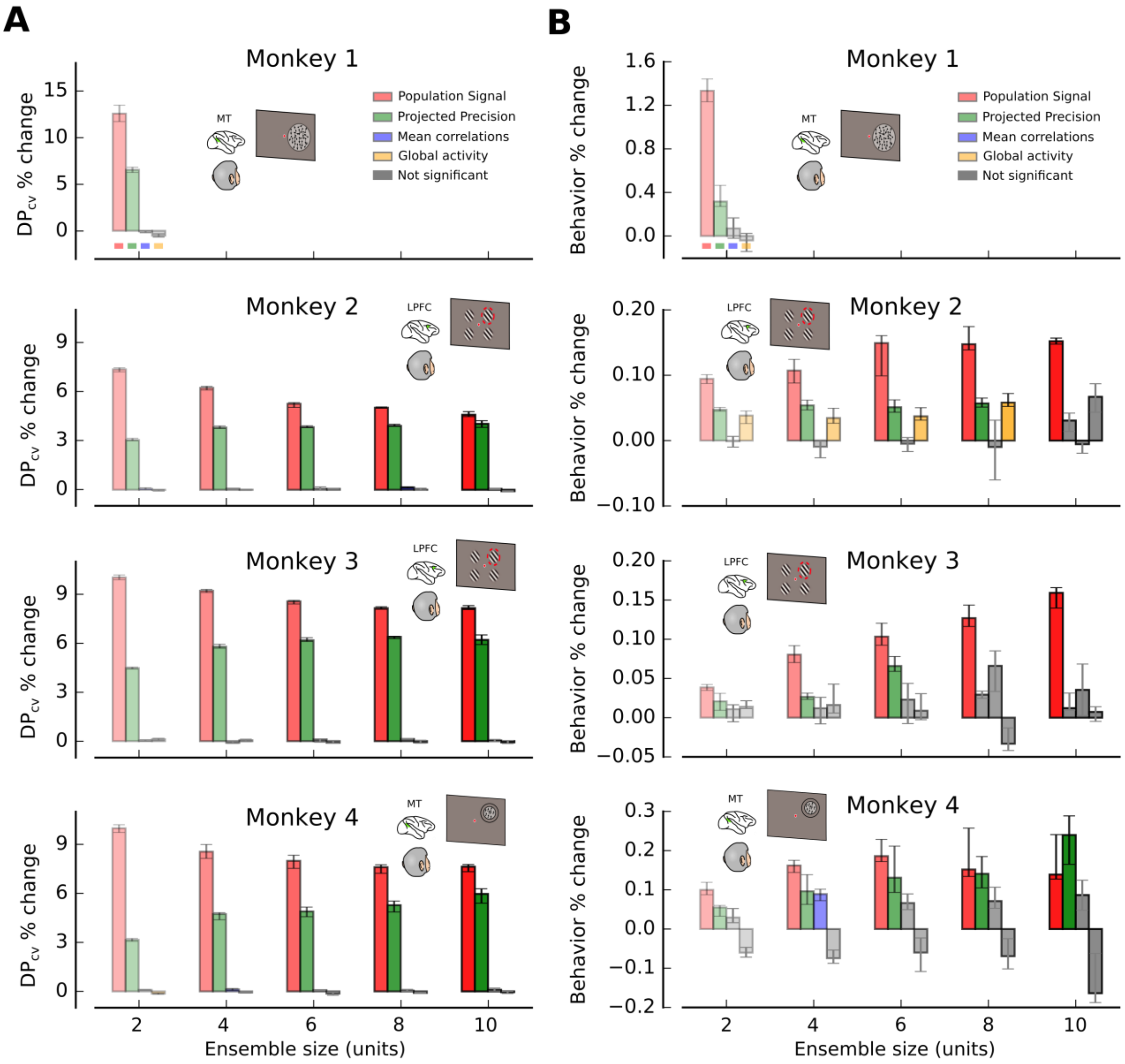


**Figure S8. Conditioning to global activity (GA) instead of mean pairwise correlations (MPC) produces equivalent results to Figs. 5 and 7. Related to Figs. 5 and 7.**

**(A)** Bootstrap fluctuations of population signal (PS) and projected precision (PP) have the largest effect on encoded information ($DP_{cv}$) when conditioning to GA (equivalent to Fig. 5). **(B)** Bootstrap fluctuations of PS and PP have the largest effect on behavioral performance when conditioning to GA (equivalent to Fig. 7).

| | | PS | PP | MPC | GA |
|---|---|---|---|---|---|
| **Monkey 1** | **2 units** | 5.84%*** | 3.21%*** | 0.00% | 0.00% |
| **Monkey 2** | **2 units** | 7.40%*** | 3.07%*** | 0.06%* | -0.02% |
| | **4 units** | 6.21%*** | 3.64%*** | 0.06% | 0.003% |
| | **6 units** | 5.31%*** | 3.85%*** | 0.05% | 0.06% |
| | **8 units** | 5.02%*** | 4.09%*** | 0.15%** | 0.002% |
| | **10 units** | 4.78%*** | 3.89%*** | 0.06% | -0.09% |
| **Monkey 3** | **2 units** | 9.90%*** | 4.51%*** | 0.04% | 0.12% |
| | **4 units** | 9.29%*** | 5.73%*** | -0.10% | 0.05% |
| | **6 units** | 8.67%*** | 6.19%*** | 0.11% | -0.10% |
| | **8 units** | 8.19%*** | 6.50%*** | 0.13% | -0.07% |
| | **10 units** | 7.94%*** | 6.53%*** | 0.03% | -0.07% |
| **Monkey 4** | **2 units** | 8.37%*** | 2.85%*** | 0.03% | -0.08%* |
| | **4 units** | 7.20%*** | 3.58%*** | 0.06% | -0.02% |
| | **6 units** | 7.06%*** | 3.89%*** | 0.07% | -0.01% |
| | **8 units** | 6.76%*** | 4.48%*** | 0.03% | -0.05% |
| | **10 units** | 6.27%*** | 4.55%*** | 0.08% | 0.007% |

**Table S1.** Percentage change on the amount of information encoded by the neuronal population (% change $DP_{cv}$) for all monkeys and ensemble sizes. Population signal and projected precision are the most influential factors on $DP_{cv}$. * = $0.01 < P < 0.05$, ** = $0.001 < P < 0.01$, *** = $P < 0.001$. Related to Fig. 5.

| | | PS - PP | PS - MPC | PS - GA | PP - MPC | PP - GA | MPC - GA |
|---|---|---|---|---|---|---|---|
| **Monkey 1** | **2 units** | 0.76%*** | 5.87%*** | 5.71%*** | 3.17%*** | 3.25%*** | 0.00% |
| **Monkey 2** | **2 units** | 4.50%*** | 7.23%*** | 7.49%*** | 3.01%*** | 3.08%*** | 0.15%* |
| | **4 units** | 2.36%*** | 6.10%*** | 6.22%*** | 3.56%*** | 3.77%*** | 0.05% |
| | **6 units** | 1.40%*** | 5.10%*** | 5.28%*** | 3.69%*** | 3.73%*** | 0.11% |
| | **8 units** | 0.91%*** | 4.90%*** | 4.67%*** | 3.80%*** | 3.87%*** | 0.11% |
| | **10 units** | 0.81%*** | 4.75%*** | 4.79%*** | 3.81%*** | 3.96%*** | 0.10% |
| **Monkey 3** | **2 units** | 5.57%*** | 1.00%*** | 1.00%*** | 4.56%*** | 4.32%*** | -0.08% |
| | **4 units** | 3.50%*** | 9.30%*** | 9.01%*** | 5.88%*** | 5.45%*** | -0.05% |
| | **6 units** | 2.40%*** | 8.29%*** | 8.74%*** | 5.95%*** | 6.33%*** | 0.08% |
| | **8 units** | 1.31%*** | 8.02%*** | 8.21%*** | 6.58%*** | 6.57%*** | 0.07% |
| | **10 units** | 1.71%*** | 7.86%*** | 8.06%*** | 6.38%*** | 6.58%*** | 0.11% |

| | | | | | | | |
|---|---|---|---|---|---|---|---|
| **Monkey 4** | **2 units** | 5.35%*** | 8.31%*** | 8.32%*** | 3.00%*** | 3.06%*** | 0.001% |
| | **4 units** | 3.65%*** | 7.74%*** | 7.71%*** | 3.69%*** | 3.71%*** | 0.13% |
| | **6 units** | 2.62%** | 7.40%** | 7.38%** | 4.27%** | 4.11%** | 0.21% |
| | **8 units** | 1.92%** | 7.42%** | 7.19%** | 4.60%** | 4.74%** | 0.07% |
| | **10 units** | 1.60%* | 6.89%* | 6.88%* | 4.67%* | 4.77%* | 0.15% |

**Table S2.** Difference of percentage change on the amount of information encoded by the neuronal population (% change $DP_{cv}$) for all monkeys and ensemble sizes for all the pairs of studied features. Population signal and projected precision are the most influential factors on $DP_{cv}$. * = $0.01 < P < 0.05$, ** = $0.001 < P < 0.01$, *** = $P < 0.001$. Related to Fig. 5.

| | | PS | PP | MPC | GA |
|---|---|---|---|---|---|
| **Monkey 1** | **2 units** | 1.41%*** | 0.56%** | 0.07% | -0.04% |
| **Monkey 2** | **2 units** | 0.114%*** | 0.041%*** | -0.001% | 0.038%** |
| | **4 units** | 0.115%*** | 0.069%*** | -0.010% | 0.035%* |
| | **6 units** | 0.138%*** | 0.076%*** | -0.005% | 0.037%* |
| | **8 units** | 0.166%*** | 0.062%** | -0.010% | 0.058%** |
| | **10 units** | 0.173%** | 0.047%* | -0.005%** | 0.067% |
| **Monkey 3** | **2 units** | 0.037%*** | 0.025%** | 0.011% | 0.014% |
| | **4 units** | 0.070%*** | 0.031%** | 0.012% | 0.016% |
| | **6 units** | 0.090%*** | 0.046%*** | 0.023% | 0.009% |
| | **8 units** | 0.074%*** | 0.036%** | 0.066% | -0.033% |
| | **10 units** | 0.153%** | 0.056%** | 0.035% | 0.007% |
| **Monkey 4** | **2 units** | 0.120%*** | 0.066%*** | 0.029% | -0.060% |
| | **4 units** | 0.138%*** | 0.008%** | 0.089%* | -0.074% |
| | **6 units** | 0.198%*** | 0.142%*** | 0.066% | -0.060% |
| | **8 units** | 0.180%** | 0.147%** | 0.071% | -0.069% |
| | **10 units** | 0.201%** | 0.182%* | 0.087% | -0.165% |

**Table S3.** Percentage change on behavioral performance (% change Behavior) for all monkeys and ensemble sizes. Population signal and projected precision are the most influential factors on behavior. * = $0.01 < P < 0.05$, ** = $0.001 < P < 0.01$, *** = $P < 0.001$. Related to Fig. 7.

| | | PS - PP | PS - MPC | PS - GA | PP - MPC | PP - GA | MPC - GA |
|---|---|---|---|---|---|---|---|
| **Monkey 1** | **2 units** | 0.690%*** | 1.31%*** | 1.26%*** | 0.462%* | 0.587%** | 0.313% |
| **Monkey 2** | **2 units** | 0.065%*** | 0.098%*** | 0.067%*** | 0.039%*** | 0.019% | -0.026%** |
| | **4 units** | 0.067%*** | 0.109%*** | 0.104%*** | 0.069%*** | 0.010%* | -0.064%** |
| | **6 units** | 0.095%** | 0.117%*** | 0.107%*** | 0.095%** | 0.024% | -0.037%* |
| | **8 units** | 0.106%** | 0.161%*** | 0.138%** | 0.047%* | -0.031% | -0.070%** |
| | **10 units** | 0.106%* | 0.192%** | 0.120%* | 0.052% | 0.018% | -0.045% |
| **Monkey 3** | **2 units** | 0.006% | 0.043%*** | 0.017% | 0.015% | 0.010% | -0.007% |
| | **4 units** | 0.062%** | 0.060%** | 0.064%* | 0.006% | -0.003% | -0.000% |
| | **6 units** | 0.023% | 0.087%** | 0.033%* | 0.047% | 0.024% | -0.007% |
| | **8 units** | 0.052% | 0.041% | 0.127%* | -0.019% | 0.077%* | 0.088% |
| | **10 units** | 0.131%* | 0.126%** | 0.156%* | 0.033% | 0.060% | 0.040% |

| | | | | | | | |
|---|---|---|---|---|---|---|---|
| **Monkey 4** | **2 units** | 0.047%** | 0.115%*** | 0.201%*** | 0.061%** | 0.103%*** | 0.090%* |
| | **4 units** | 0.118%* | 0.067%** | 0.243%** | 0.021% | 0.143%** | 0.128% |
| | **6 units** | 0.080%* | 0.140%** | 0.310%** | 0.087%** | 0.178%** | 0.150% |
| | **8 units** | 0.070% | 0.170%** | 0.403%* | 0.150%** | 0.279%* | 0.141% |
| | **10 units** | 0.099% | 0.212%* | 0.412% | 0.113% | 0.160%* | 0.161% |

**Table S4.** Difference of percentage change on behavioral performance (% change Behavior) for all monkeys and ensemble sizes for all the pairs of studied features. Population signal and projected precision are the most influential factors on behavior * = $0.01 < P < 0.05$, ** = $0.001 < P < 0.01$, *** = $P < 0.001$. Related to Fig. 7.
